# Reduction of RAD23A extends lifespan and mitigates pathology in TDP-43 mice

**DOI:** 10.1101/2024.09.10.612226

**Authors:** Guo Xueshui, Ravindra Prajapati, Jiyeon Chun, Insuk Byun, Kamil K Gebis, Yi-Zhi Wang, Karen Ling, Casey Dalton, Jeff A. Blair, Anahid Hamidianjahromi, Gemma Bachmann, Frank Rigo, Paymaan Jafar-nejad, Jeffrey N. Savas, Min Jae Lee, Jemeen Sreedharan, Robert G. Kalb

**Author notes:** Ashfield MedComms, 300 Vesey Street, 10th Floor, New York, NY 10282.

## Abstract

Protein misfolding and aggregation are cardinal features of neurodegenerative disease (NDD) and they contribute to pathophysiology by both loss-of-function (LOF) and gain-of-function (GOF) mechanisms. This is well exemplified by TDP-43 which aggregates and mislocalizes in several NDDs. The depletion of nuclear TDP-43 leads to reduction in its normal function in RNA metabolism and the cytoplasmic accumulation of TDP-43 leads to aberrant protein homeostasis. A modifier screen found that loss of *rad23* suppressed TDP-43 pathology in invertebrate and tissue culture models. Here we show in a mouse model of TDP-43 pathology that genetic or antisense oligonucleotide (ASO)-mediated reduction in *rad23a* confers benefits on survival and behavior, histological hallmarks of disease and reduction of mislocalized and aggregated TDP-43. This results in improved function of the ubiquitin-proteasome system (UPS) and correction of transcriptomic alterations evoked by pathologic TDP-43. RAD23A-dependent remodeling of the insoluble proteome appears to be a key event driving pathology in this model. As TDP-43 pathology is prevalent in both familial and sporadic NDD, targeting *RAD23A* may have therapeutic potential.

## Introduction

Cytoplasmic, nuclear and neuritic ubiquitinated inclusions (UBI) are a pathological hallmark of amyotrophic lateral sclerosis (ALS), frontotemporal lobar dementia (FTD) and subtypes of other NDD^1^. TAR-DNA-binding protein-43 (TDP-43) was originally identified as a disease-specific core protein of a subpopulation of UBIs^2^. We now know that TDP-43 is a critical disease driver due to both LOF and GOF mechanisms that are linked to its mislocalization. Strategies to ameliorate pathophysiological events evoked by TDP-43 are likely to have a major therapeutic impact on ALS/FTD.

Unbiased screens for phenotypic suppressors evoked by overexpression of wild type (WT) or mutated versions of TDP-43 have shown that many biological processes are impacted including components of the proteostasis network^3–10^. It has been estimated that 80% of cellular proteins are degraded by UPS^11^. Ubiquitinated clients can bind to the proteasome (i.e., Rpn1/PSMD2, Rpn10/PSMD4 and Rpn13/ADRM1)^12^ to initiate proteolysis by the 20S proteasome. An additional route of those clients to the proteasome employs “shuttle factors” such as yeast proteins RAD23 (RAD23A and RAD23B in vertebrates), Dsk2 (ubiquilins, UBQLNs 1 – 4 in vertebrates) and Ddi1 (Ddi1 and Ddi2 in vertebrates)^13^. While the canonical view of RAD23 function as a shuttle factor that promotes client degradation is clearly correct for many substrates^14–18^, it is not the complete story. Loss of rad23 can also promote the degradation of targets such as XPC^19^, p53^20^, Sic1^16^, polyQ expanded ataxin 3^21^, mutant SOD and mutant TDP-43^22^. This may be linked to features of: 1) the substrate, as well-folded clients brought to the proteasome by RAD23 resist degradation^23^ or 2) ubiquitin status, as RAD23 negatively regulates multi-ubiquitin chain assembly^24,25^.

Ablation of RAD23 in a *Caenorhabditis elegans* model of TDP-43’opathy conferred benefits on locomotor and biochemical phenotypes and as does knockdown (KD) of the two vertebrate isoforms of RAD23 (RAD23A or RAD23B) in a rat neuron tissue culture model^22^. Mechanistic insights into RAD23A/B actions in a TDP-43-based mouse model of ALS/FTD might empower therapeutic opportunities.

## Results

Potent and specific ASOs were designed targeting *RAD23A* or *RAD23B* mRNA. After intracerebroventricular (ICV) injection at postnatal day zero (P0) of mouse development of these ASOs (or a non-targeting control sequence, scrambled or “SCR”), we assessed the effects on *RAD23A* and *RAD23B* mRNA levels. In the cerebral cortex, *RAD23A* mRNA level was reduced ∼65% (ASO: RAD23A versus SCR, p=0.0001) and in the spinal cord reduced ∼70% (RAD23A versus SCR, p=0.0001)(Figure 1A). The ASO targeting *RAD23A* had no effect on the *RAD23B* mRNA level in the cerebral cortex or spinal cord (RAD23A versus SCR, p=0.71; RAD23A versus SCR, p=0.11, respectively Figure 1B). The ASO targeting *RAD23A* mRNA led to an ∼80% KD of brain RAD23A protein (RAD23A versus SCR, p=0.0001, Figure 1C,D) and no effect on RAD23B protein level (RAD23A versus SCR, p=0.89) Figure 1C,D). We next turned to the TAR4/4 mouse that expresses WT human TDP-43 (hTDP-43) under the control of the Thy-1 promoter^26^. These animals express hTDP-43 ∼2 time greater than the endogenous mouse TDP-43 (mTDP-43) and develop many histological and biochemical features seen at autopsy of individuals with ALS or FTD. Hereafter we refer to TAR4/4 mice with KD of RAD23A owing to ASOs as TAR4/4;RAD23A^KD^ and control mice receiving the scrambled sequence ASO as TAR4/4;RAD23A^SCR^. In comparison with the TAR4/4;RAD23A^SCR^ mice, the TAR4/4;RAD23A^KD^ males and females had a ∼9 day extension in life span (∼50% increase in lifespan, hazard ratio (HR) 10.15, p<0.0001) (Figure 1E). The effects in males were a ∼7 day increase in lifespan (HR=7.12, p<0.0001) and females a ∼10 day increase in lifespan (HR=29.12, p<0.0001) (Supplemental Figure 1A,B). We ran a second independent cohort of TAR4/4 mice and found an identical pro-survival effect of knockdown of RAD23A in both female and male mice (Supplemental Figure 1C-E). The TAR4/4 animals develop several abnormal motor phenotypes that worsen over time and KD of RAD23A slowed the development of the gait impairment, tremor and kyphosis phenotypes and hindlimb clasping (Figure 1F-I, Supplemental Figure 1F).

**Figure 1.**
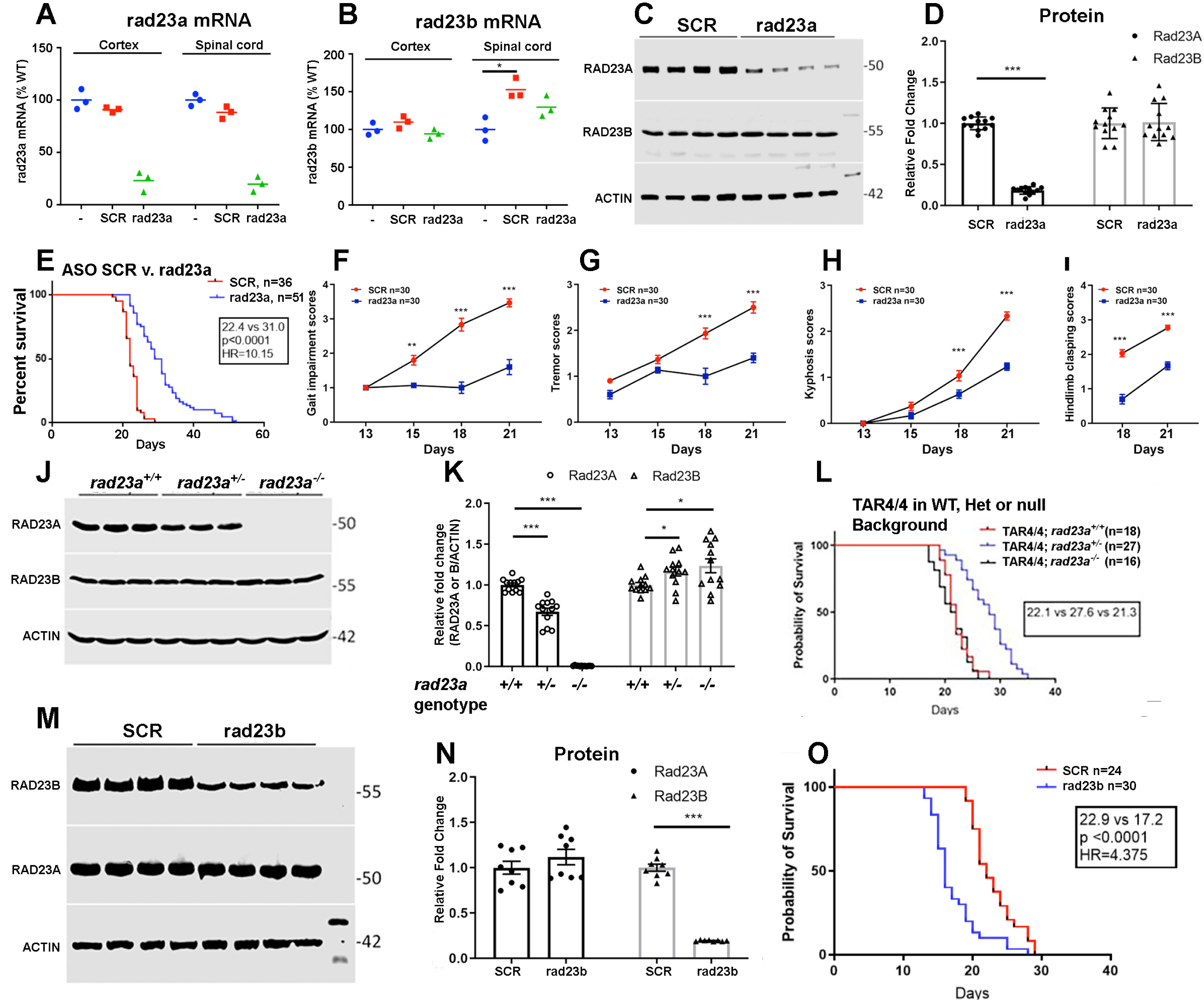
Effect of RAD23A KD on TAR4/4 mice: survival and behavior. (A) Quantification of the ASO effect on *RAD23A* mRNA level. Group differences were found by ANOVA F_(2,6)_= 83.63, p<0.0001, post hoc analysis: ***, p<0.001 (B) Quantification of the ASO effect on *RAD23B* mRNA level. Group differences were found by ANOVA F_(2,6)_= 3.26, p<0.01, post hoc analysis: ***, p,0.001 (C) Western blots of cortex from mice that received intracerebroventricular (ICV) administration at postnatal day zero (P0) of an antisense oligonucleotide (ASO) with a scrambled sequence (SCR) or targeting rat *RAD23A.* ASO to *RAD23A* leads to a reliable and reproducible reduction in abundance of RAD23A, but not RAD23B, protein. N=4 shown. (D) Quantification of the ASO effects at the protein level. ***, p<0.001. (E) Survival of TAR4/4 male + female mice after administration of ASO targeting *RAD23A* or the SCR control. There is an ∼9 day extension of lifespan in animals receiving the ASO to *RAD23A*. The lifespan curves were compared by the log-rank test (<0.0001) and the effect size was estimated by the Cox proportional hazards model, where HR indicated the hazard ratio (HR=10.15). (F) Progressive rise in the gait impairment score in the TAR4/4 mice treated with SCR ASO was blunted by ASO targeting RAD23A. **, p<0.01 and ***, p<0.001. (G) Progressive rise in the tremor score in the TAR4/4 mice treated with SCR ASO was blunted by ASO targeting *RAD23A*. (H) Progressive rise in the kyphosis score in the TAR4/4 mice treated with SCR ASO was blunted by ASO targeting *RAD23A*. ***, p<0.001. (I) Progressive rise in the hindlimb clasping score in the TAR4/4 mice treated with SCR ASO was blunted by ASO targeting *RAD23A*. ***, p<0.001. ***. Note, hindlimb clasping cannot be reliably assessed prior to P18. (J) There is a reduction of RAD23A (but not RAD23B) protein levels in mice heterozygous for the RAD23A null allele (*RAD23A^-/+^*) and no RAD23A expression in the homozygotes (RAD23A^-/-^). (K) Quantification of genotype effect on RAD23A and RAD23B protein levels. Group differences were found by ANOVA for RAD23A - F(2,33)= 306.2, p<0.0001, post hoc analysis: *, p<0.05, ***, p<0.001. Group differences were found by ANOVA for RAD23B - F_(2,33)_= 3.90, p<0.03, post hoc analysis: *, p<0.05, ***, p<0.001. (L) Survival of TAR4/4 mice in the background of WT, reduced or absent RAD23A levels. There is an ∼5 day extension of lifespan in TAR4/4;RAD23A^-/+^ animals in comparison with TAR4/4;*RAD23A^+/+^*or TAR4/4;RAD23A^-/-^ animals (HR=9.4, p<0.001 TAR4/4;*RAD23A^-/+^*versus TAR4/4;RAD23A^+/+^). Number of animals in experimental groups (TAR4/4;RAD23A^+/+^, TAR4/4;RAD23A^-/+^ and TAR4/4;RAD23A^-/-^) was 18,27 and 16 respectively. (M) Western blots of cortex from mice that received SCR ASO or ASO targeting *RAD23B* leads to a reliable and reproducible reduction in abundance of RAD23B but not RAD23A protein. N=4 shown. (N) Quantification of the ASO effects at the protein level. ***, p<0.001. (O) Survival of TAR4/4 mice treated with SCR or *RAD23B* targeting ASO. There is an ∼5 day shortening of lifespan of animals associated with RAD23B protein KD extension (HR=4.3, p<0.001).

We wondered if complete elimination of RAD23A would have an even more robust pro-survival effect on the TAR4/4 mouse. We generated mice with a RAD23A null allele using CRISPR/Cas9; RAD23A^-/-^ have no detectable RAD23A protein while RAD23A^+/-^ animals have an ∼35% reduction in RAD23A protein (WT versus RAD23A^+/-^, p<0.0001) (Figure 1J,K). In comparison with WT animals, there is a ∼15% and ∼20% increase in RAD23B protein in the RAD23A^+/-^ and RAD23A^-/-^ mice, respectively (p<0.05 for both, Figure 1K). To our surprise, there is no lifespan extension in the TAR4/4;RAD23A^-/-^ mice. TAR4/4;RAD23A^+/-^ display an ∼5 days increase in lifespan (compared with TAR4/4;RAD23A^+/+^ HR=9.49, p<0.0001) (Figure 1L). The absence of a life span extension phenotype in the TAR4/4;RAD23A^-/-^ mice may indicate that compensatory changes evoked by germline loss of RAD23A confound any beneficial effects on the TAR4/4 disease or that some residual level of RAD23A protein is necessary the beneficial effects on the TAR4/4 phenotypes.

We developed an ASO targeting RAD23B and in comparison to the SCR ASO, it led to a ∼90% reduction in RAD23B protein (SCR versus RAD23B, p=0.0001) and had no effect on RAD23A protein (SCR versus RAD23A, p=0.30) (Figure 1M,N). In comparison with the SCR ASO delivered ICV at identical concentration, KD of RAD23B led to an ∼5 day shortening of lifespan (HR=4.37, p<0.0001) (Figure 1O).

As reported, the TAR4/4 motor cortex displays reduced thickness and neuronal density^26^. We saw a reduction in motor cortex NeuN(+) neurons associated with increased number of glial fibrillary astrocytic protein positive (GFAP+) cells predominantly in layers II and V (Figure 2A). TAR4/4;RAD23A^KD^ mice have a thicker motor cortex with a higher density of NeuN(+) neurons and less astrocytosis in comparison with TAR4/4;RAD23A^SCR^ mice. In both the cerebral cortex and spinal cord, there is an increase in the GFAP transcript in the TAR4/4;RAD23A^SCR^ mice in comparison with WT animals that is reduced in the TAR4/4;RAD23A^KD^ mice (Figure 2B). Similar effects were seen with two microglial markers IBA1 and CD68 (Figure 2C,D). We find cleaved caspase 3 in the cerebral cortex of TAR4/4;RAD23A^SCR^ mice (and not in WT mice) and there is ∼50% decrease (p<0.001) in the abundance of this species in the TAR4/4;RAD23A^KD^ mice. The level of uncleaved caspase 3 is modestly elevated in TAR4/4;RAD23A^SCR^ versus TAR4/4;RAD23A^KD^ (∼15%, p<0.05) (Figure 2E-G). Together these observations reveal neuron death along with astro- and micro-gliosis in the cerebral cortex of the TAR4/4 mice and these pathologies are blunted by KD of RAD23A.

**Figure 2.**
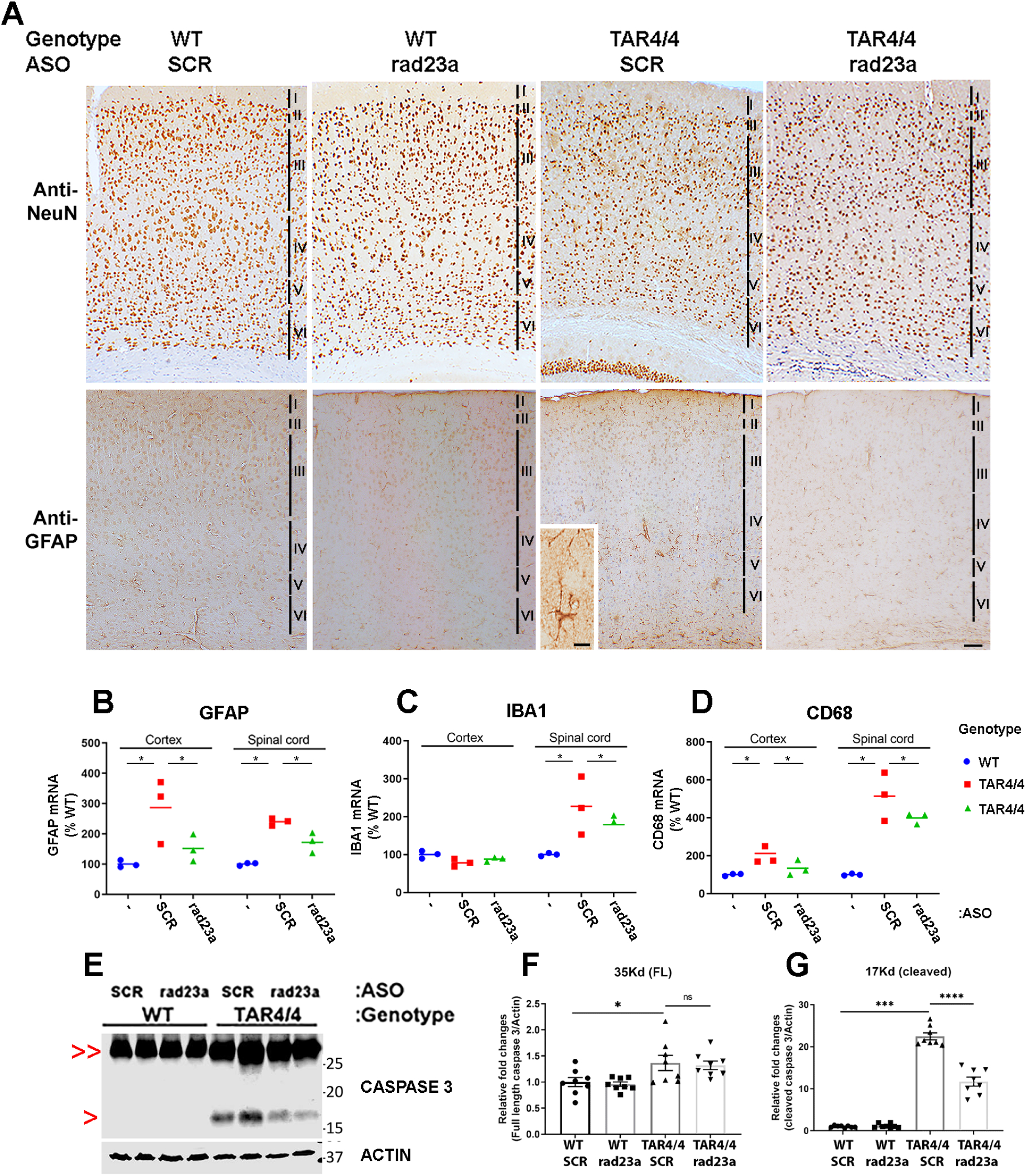
Effect of RAD23A on TAR4/4 mice: histology and cell type markers. (A) Immunohistology for a marker of neuronal nuclei (anti-NeuN) and astrocytes (anti-glial fibrillary acidic protein, GFAP) in WT and TAR4/4 mice who received ICV injection at P0 of SCR sequence or RAD23A targeted ASO. To study primary and secondary motor cortex, coronal tissue slices correspond to Bregma ∼1.3 mm (assuming interaural position at 2.48 mm). At this level, a portion of hippocampus (e.g., CA1, CA2, CA3 and dentate gyrus) is visible ventrally. The motor cortex of WT animals treated with ASO SCR or targeting RAD23A display indistinguishable thickness and neuronal density. The same is true for GFAP immunoreactivity. The motor cortex of TAR4/4 animals treated with the ASO SCR is less thick and displays a reduced density of neurons particularly in layers VI and V. These layers contain a prominent population of GFAP(+) astrocytes. Astrocytes also populate more superficial layers (particularly layer II). Blow up of astrocyte, inset. The motor cortex of TAR4/4 animals treated with the ASO to *RAD23A* blunt these pathological findings with partial restoration of cortical thickness and density and reduced astrocytosis. Calibration bar of high lower inset = 15 microns. Calibration bar from low power octet = 50 microns (B – D) Quantitative PCR for GFAP (B), IBA1 (C), and CD68 (D) in WT animals and TAR4/4 animals that received ICV injection of ASO at P0 that are SCR sequence or target *RAD23A*. (B) SCR treated TAR4/4 animals have an elevated level of GFAP transcripts in comparison with WT animals and this is blunted in TAR4/4 animals treated with ASO targeting *RAD23A*. This is seen in both the cerebral cortex and spinal cord (p < 0.05). (C) There are no group difference in the cortical IBA1 transcripts. In the spinal cord, IBA1 transcripts are elevated in the SCR treated TAR4/4 animals and this is blunted in TAR4/4 animals treated with ASO targeting *RAD23A* (p < 0.05). SCR treated TAR4/4 animals have a modestly elevated level of CD68 transcripts in comparison with WT animals and this is blunted in TAR4/4 animals treated with ASO targeting *RAD23A*. This is seen in both the cerebral cortex and spinal cord (p > 0.05). (E) Full length (FL) caspase 3 is detectable in duplicate WT and TAR4/4 animals and its level of abundance is higher in the TAR4/4 animals regardless of ASO administered (denoted by red >> symbol). Only in the TAR4/4 animals is the cleaved caspase 3 species detectable (denoted by red >) and the level of abundance is higher in the SCR ASO treated animals in comparison with the RAD23A ASO treated animals. (F) Quantification of the abundance of FL caspase 3 reveals a ∼30% increase in the TAR4/4 compared with WT mice (p<0.05). (G) Quantification of the abundance of cleaved caspase 3 reveals a ∼50% decrease in the TAR4/4 RAD23A ASO treated compared with SCR ASO treated mice (p < 0.0001).

Normally, TDP-43 is observed almost exclusively in the nucleus and pathologic depletion of nuclear TDP-43 causes RNA processing LOF phenotypes. Since mislocalization of TDP-43 has been reported in the TAR4/4 mice^26^ we wondered if this affected RNA biology and a potential relationship to rad23a. To this end we undertook a transcriptomic interrogation of the cortex of four groups of animals: 1) Group I - TAR4/4;RAD23A^SCR^ versus WT;RAD23A^SCR^ revealed 14,312 differentially expressed genes (DEGs; 7658 up, 6651 down) representing the “disease genes” (Supplemental material Figure 2), and 2) Group II - TAR4/4;RAD23A^KD^ versus TAR4/4;RAD23A^SCR^ revealed 8,108 DEGs (4,022 up, 4,486 down) representing “treatment genes” (Supplemental material Figure 3, Figure 3A,B). Rad23a displayed the lowest adjusted P-value (3.99E-48) and lowest log2 fold-change (-1.6) in the Group 2 DEGs comparison, confirming its successful knockdown (Figure 3B). 6,125 DEGs show reversible expression patterns (upregulated in TAR4/4;RAD23A^SCR^ were downregulated in TAR4/4;RAD23A^KD^ and down-regulated in TAR4/4;RAD23A^SCR^ were upregulated in TAR4/4;RAD23A^KD^), providing evidence that RAD23A KD significantly blunts the effects of hTDP43 on the transcriptome. This is well illustrated by the principal component analysis and unsupervised hierarchical clustering heatmap. These analysis reveal that experimental groups are distinct both in terms of up and down regulated transcripts (Figure 3C,D) and reversable in a RAD23A-dependent manner. We assessed changes in the abundance of 15 transcripts by qPCR and validated the RNAseq results through this independent method (Supplemental Figure 4)

**Figure 3.**
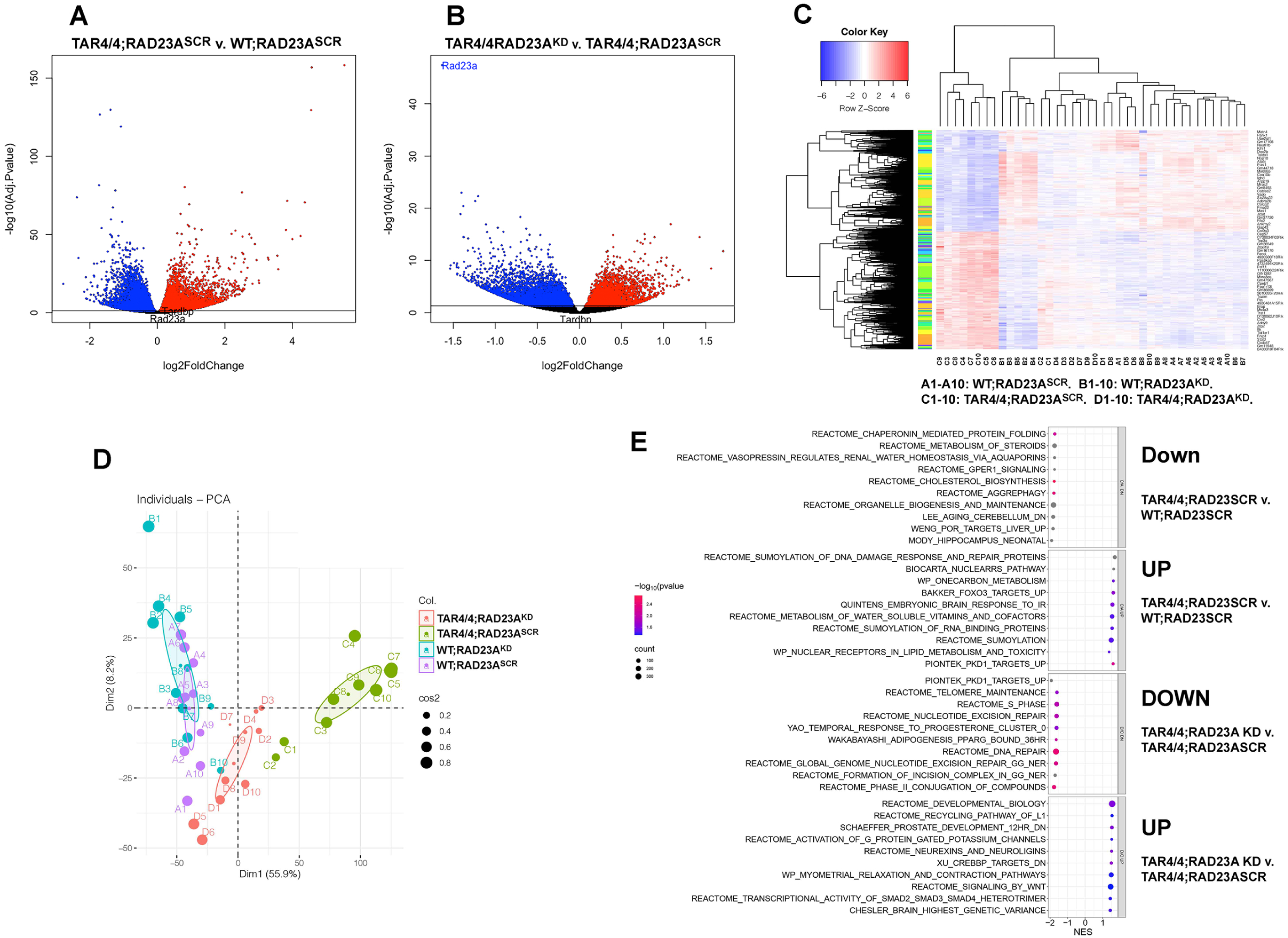
Effects on hTDP-43 on the mouse transcriptome are modified by KD of RAD23A. (A) The volcano plots represent the differentially expressed genes (DEGs) in the pairwise comparisons between TAR4/4;RAD23A^SCR^ versus WT;RAD23A^SCR^. Genes with an adjusted P-value of less than 0.05 were considered as DEGs. Upregulated and downregulated DEGs shown in red and blue, respectively. Genes with a positive log2(foldchange) are considered upregulated, while those with a negative log2(foldchange) are considered downregulated. The horizontal line represents the adjusted P-value cut-off (0.05). (B) The volcano plots represent the DEGs in the pairwise comparisons between TAR4/4;RAD23A^SCR^ versus TAR4/4;RAD23^KD^. Statistical considerations as in (A). (C) Hierarchical clustering and heatmap generation was performed using 6121 genes which show a reversed expression patterns between TAR4/4 and TAR4/4 ASOs(RAD23A). All four conditions, WT and TAR4/4 mice treated with SCR ASO or ASO targeting *RAD23A* (10 replicates). The KD of RAD23A in the TAR4/4 mice leads to a shift in the transcriptomes towards the WT. The color key indicates the row Z-score (red: upregulation after treatment with RAD23A ASOs; blue: downregulation with RAD23A treatment). (D) Principal Component Analysis (PCA) was performed on 6121 genes that displayed reversed expression patterns between TAR4/4 SCR ASO versus TAR4/4 RAD23A targeting ASO. All four experimental groups and all their replicates were included in the PCA. FactoMineR and Factoextra R-packages were used for performing the PCA analysis. (E) Gene ontology terms revealing reversible enrichment patterns between the experimental groups. Pairwise comparisons of the Group I comparisons (Up regulated and down regulated transcripts are displayed separately); similarly Group II comparisons. The circle size corresponds to the number of genes, and the blue-to-red colour gradient represents the P-value.

A Gene Set Enrichment Analysis (GSEA) focusing on the top ten up and down-regulated gene ontology (GO) terms (Figure 3E) reveals that in comparison with WT;RAD23A^SCR^, the TAR4/4;RAD23A^SCR^ mice exhibit upregulation of sumoylation in DNA damage response and RNA binding proteins and downregulation of aggrephagy and chaperone-mediated protein folding. A reduction in the transcription of protein homeostasis machinery likely contributes to the accumulation of aggregated proteins. In comparison with the TAR4/4;RAD23A^SCR^, the TAR4/4;RAD23A^KD^ mice exhibit upregulation of neurexins and neuroligins, G-protein gated potassium channels, Wnt signaling, and Smad2, 3, & 4 heterotrimer signalling and down regulation of DNA repair, nucleotide excision repair, and telomere maintenance (Figure 3E). This pattern of up-regulated gene expression may indicate that KD of rad23 blunts disruption (or promotes repairs) in synapse formation and signaling evoked in TAR4/4;RAD23A^SCR^ mice. Down-regulation of DNA repair process may indicate that KD of rad23a improves genomic integrity in TAR4/4 mice and/or that Rad23A is required for the expression of DNA repair genes.

We next identified reversibly enriched GO terms between Group 1 and Group 2, and found 25 gene ontology exhibiting reversed enrichment (Supplemental Figures 5,6) GO terms associated with neural function, neural development, mitochondrial function, and mitochondria biogenesis exhibit reversible enrichment patterns; down-regulated in the TAR4/4;RAD23A^SCR^ in comparison to WT;RAD23A^SCR^ and upon Rad23A KD, they were upregulated. In addition, aggrephagy (but not sumoylation) was down regulated in the TAR4/4;rad23a^SCR^ (in comparison with WT;RAD23A^SCR^) and also observed upregulation of aggrephagy in TAR4/4;rad23a^KD^ and upon Rad23A KD, there was upregulation of aggrephagy genes. This presumably creates favorable conditions for clearing the protein aggregate from the cells and promote a healthy proteostasis network.

GOF phenotypes evoked by TDP-43 are dose-dependent^26^ and so we wondered whether the beneficial effects of RAD23A KD influenced TDP-43 abundance. There is ∼50% reduction of hTDP-43 expression in the TAR4/4;RAD23A^KD^ versus TAR4/4;RAD23A^SCR^ brain (p<0.01) as well as 50% reduction of mouse plus human TDP-43 (m+hTDP-43) expression in the brain (p<0.01) (Figure 4A-C). We did not detect a difference in the abundance of the hTDP-43 transcript in TAR4/4;RAD23A^KD^ versus TAR4/4;RAD23A^SCR^ mice (cortex or spinal cord, p=0.14 and 0.69, respectively) (Figure 4D). To examine the possibility that alterations in TDP-43 abundance are compartment-specific, we generated nuclear and cytoplasmic fractions and immunoblotted for hTDP-43. In comparison with SCR ASO treatment, knockdown of *RAD23A* leads to an ∼ 70% increase in the nuclear/cytoplasmic ratio of hTDP-43 (Figure 4E; determinations of cytosolic and nuclear TDP-43 as a function of treatment are in Supplemental Figure 7A-C) and thus provides a potential mechanism by which transcriptomic derangement in the TAR4/4 brain is blunted by RAD23A KD.

**Figure 4.**
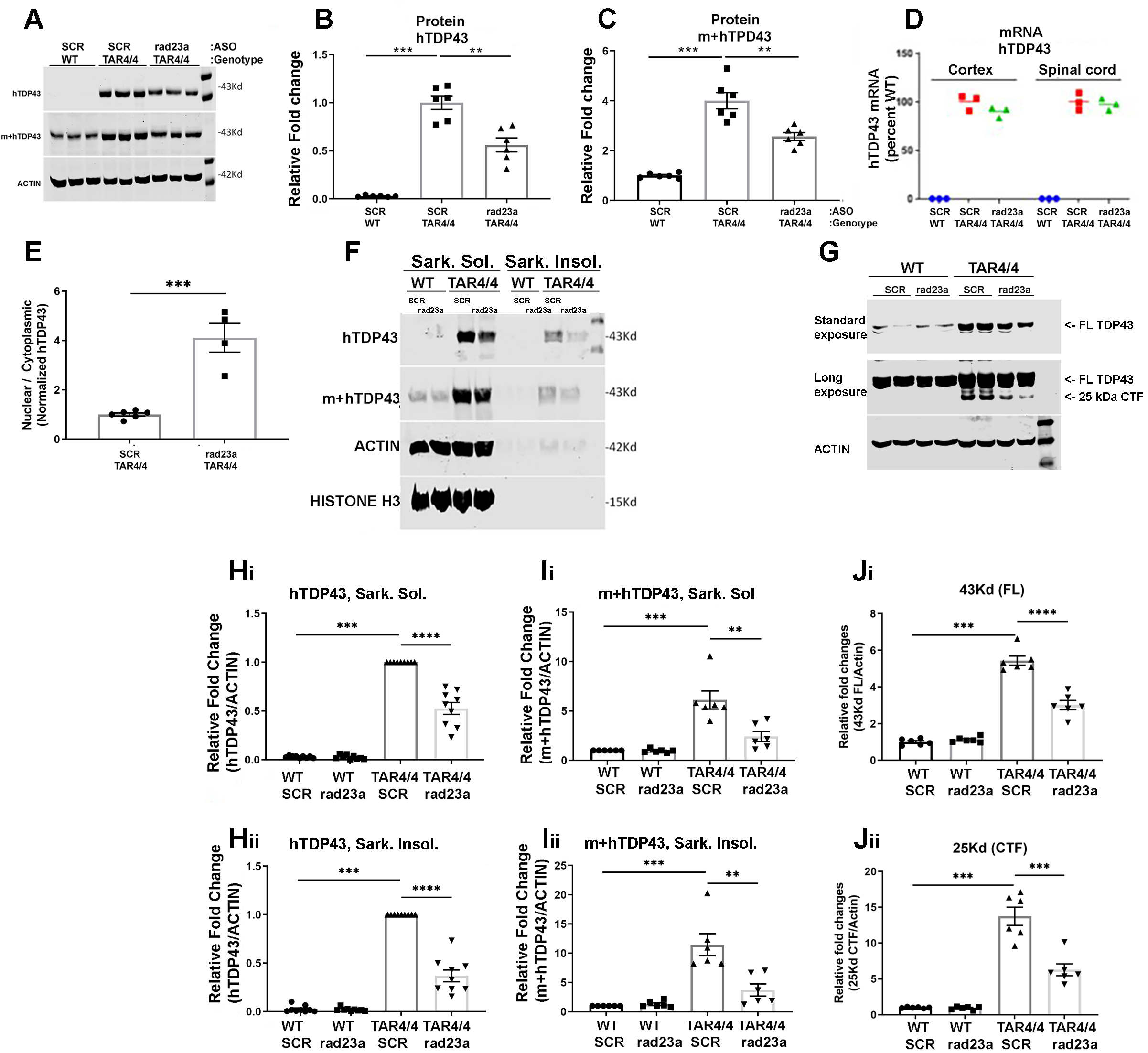
Effect of RAD23A on TAR4/4 mice: subcellular distribution and solubility of TDP-43. (A) Western blots for human TDP-43 (hTDP-43) or both mouse and human TDP-43 (m+hTDP-43) of cortex from WT and TAR4/4 mice treated with ASOs. hTDP-43 is only detectable in the cortex of TAR4/4 mice and knockdown of RAD23A reduces the abundance of TDP-43 when blotting for hTDP-43 or m+hTDP-43. (B) Quantification of the ASO effects on the hTDP-43 protein level. Group differences were found by ANOVA, F_(2,15)_= 70.5, p<0.0001, post hoc analysis: *, p<0.05, ***, p<0.001. **, p<0.01, ***, p<0.001 (C) Quantification of the ASO effects on the m+hTDP-43 protein level. Group differences were found by ANOVA, F_(2,15)_= 51.6, p<0.0001, post hoc analysis:**, p<0.01, ***, p<0.001 (D) Transcript for hTDP-43 is present only in TAR4/4 mice; there is no different in this transcript when comparing SCR ASO versus RAD23A targeting ASO in either the cortex (p=0.14) or spinal cord (p=0.69) (E) Nuclear and cytosolic fractionation of cortex from WT and TAR4/4 was performed on mice that received the SCR sequence or RAD23A targeting ASO. After normalization, the ratio of nuclear to cytosolic hTDP43 was obtained on a per experimental animal basis. There is a statistically significant increase in the nuclear/cytosolic ratio in the RAD23A ASO treated animals in comparison with the SCR ASO treate animals. ***, p<0.001 (F) Brain lysates were subjected to sarkospin protocol and separated into sarkosyl soluble (sark. sol.) and sarkosyl insoluble (sark. insol.) fractions. Human TDP-43 is enriched in the sark. sol. fraction of TAR4/4 cortex (in comparison with the sark. insol. fraction) and RAD23A ASO reduces sark. sol. and insoluble hTDP-43 in comparison with SCR treated animals. This is also true when blotting for m+hTDP-43. Actin and histone H3 are enriched in the sark. sol. fraction and not present in the sark. insol. fraction, confirming the effectiveness of the sarkospin protocol. (G) Lysates of cortex from WT and TAR4/4 animals probed for m+hTDP-43 reveal more total TDP-43 in the TAR4/4 mice. In comparison with SCR ASO, the ASO targeting RAD23A reduces full length (FL) TDP-43 (as seen in panels A and F) upon standard exposure time on the LICOR scanner. With long exposure, the 25 kDa C-terminal fragment (CTF) is detectable in TAR4/4 brain lysates and in comparison SCR ASO treated animals, the ASO targeting RAD23A reduces the abundance of the CTF. (H_i_) Quantification of the ASO effects on hTDP-43 in the sark. sol. fraction. Group differences were found by ANOVA, F_(3,32)_= 228.3, p<0.0001, post hoc analysis: ***, p<0.001, ****, p<0.0001. (H_ii_) Quantification of the ASO effects on hTDP-43 in the sark. insol. fraction. Group differences were found by ANOVA, F_(3,32)_= 221.8, p<0.0001, post hoc analysis: ***, p<0.001, ****, p<0.0001. (I_i_) Quantification of the ASO effects on m+hTDP-43 in the sark. sol. fraction. Group differences were found by ANOVA, F_(3,20)_= 21.03, p<0.0001, post hoc analysis: **, p<0.01, ***, p<0.001. (I_ii_) Quantification of the ASO effects on m+hTDP-43 in the sark. insol. fraction. Group differences were found by ANOVA, F(3,20)= 20.31, p<0.0001, post hoc analysis: **, p<0.01, ***, p<0.001. (J_i_) Quantification of the ASO effects on FL m+hTDP-43. Group differences were found by ANOVA, F_(3,20)_= 128.5, p<0.0001, post hoc analysis: ***, p<0.001. (J_ii_) Quantification of the ASO effects on CTF. Group differences were found by ANOVA, F_(3,20)_= 64.0, p<0.0001, post hoc analysis: ***, p<0.001.

In the brains of patients with ALS/FTD, a minor fraction of TDP-43 misfolds, oligomerizes and aggregates into cytotoxic species that are enriched in a sarkosyl detergent-insoluble fraction^27^. We found hTDP-43 in both the sarkosyl-soluble and -insoluble fractions of the TAR4/4 mouse brain. KD of *RAD23A* leads to an ∼50% reduction of sarkosyl-soluble and an ∼65% reduction of sarkosyl-insoluble hTDP-43 (p<0.001 and p<0.001, respectively, Figure 4F,H_i_,H_ii_). Total TDP-43 (m+h) shows similar effects; KD of RAD23A leads to an ∼75% reduction of sarkosyl-soluble and an ∼75% reduction of sarkosyl-insoluble m+hTDP-43 (SCR versus RAD23A ASO p<0.01 and p<0.01, respectively) (Figure 4F,I_i_,I_ii_). The C-terminal 25 kDa fragment of TDP-43 is a toxic species and is found in the TAR4/4 mice. KD of RAD23A reduces the abundance of both full length TDP-43 ∼40% and the 25 kDa fragment ∼60% (TAR4/4;RAD23A^KD^ versus TAR4/4;RAD23A^SCR^, p<0.0001 and p<0.001, respectively, Figure 4G,J_i_,J_ii_). These results suggest that the benefits of RAD23A KD in the TAR4/4 mice are associated with reduced accumulation of toxic forms of hTDP-43.

Another molecular hallmark of TDP-43’opathy is the accumulation and aggregation of p62 (sequestosome 1). We find a 6x increase in p62 in the insoluble, but not soluble, fraction of TAR4/4 brains in comparison with WT animals (p<0.001 and p=0.65, respectively). There is an ∼ 55% decrease in p62 in the insoluble fraction of TAR4/4;RAD23A^KD^ versus TAR4/4;RAD23A^SCR^ (p<0.001, Supplemental Figure 8A,C,D). Immunohistochemical (IHC) examination of the motor cortex reveals that in the TAR4/4, but not WT, mice p62 is distributed throughout the soma and extends extensively into proximal dendrites, principally in neurons in layers IV and V of the cortex. Discrete intense neuronal p62 immunoreactive puncta are visible (Supplemental Figure 8B).

RAD23A has two ubiquitin-associated (UBA) domains and a ubiquitin-like (UBL) domain. The UBA domains act as ubiquitin receptors, binding ubiquitinated clients and shuttling them to the proteasome for degradation through the association of the UBL domain with the proteasome. Since the characteristic signature of pathological TDP-43 includes ubiquitinated aggregates it is possible the presence or absence of RAD23A protein influences the ubiquitome and proteasomal function in general. We find a more prominent IHC signal for ubiquitin in the cerebral cortex of TAR4/4 in comparison to WT animals, in both the nucleus and proximal dendrites (Supplemental Figure 9). By immunoblot, total ubiquitin levels are ∼8 times higher in the TAR4/4;RAD23A^SCR^ versus WT;RAD23A^SCR^ brain lysates in both the soluble as well as insoluble fractions (p<0.001 and p<0.001, respectively). KD of RAD23A reduces total ubiquitin levels ∼60% and ∼45% in the insoluble and soluble fractions (RAD23A targeting ASO versus SCR, p<0.01 and p<0.001, respectively) (Figure 5A-C). Elevated levels of total ubiquitin in the TAR4/4 mice might indicate that proteasomal degradation of ubiquitinated clients is impaired in a RAD23A-dependent manner.

**Figure 5.**
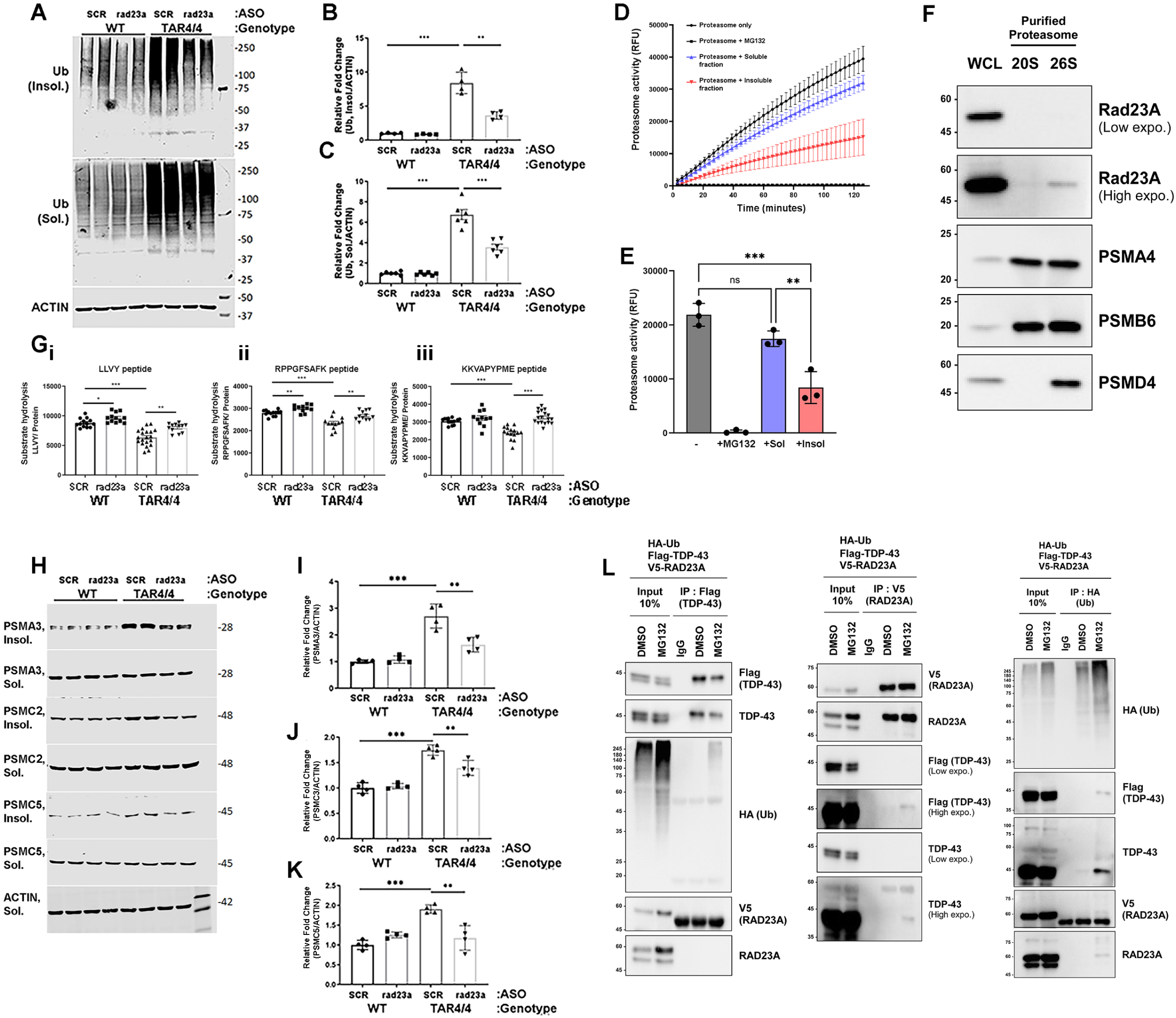
Effect of RAD23A on TAR4/4 mice: proteasome level, localization and activity. (A)Total ubiquitin levels in the soluble and insoluble fraction of WT and TAR4/4 mice treated with ASOs. The smear of ubiquitinated substrates in WT animals (insoluble and soluble fractions) is unaffected by ASO treatment. In contrast, the overall burden of total ubiquitinated substrates in the TAR4/4 mice is higher than in WT mice (in both insoluble and soluble fractions). KD of RAD23A reduces the abundance of total ubiquitinated substrates (in comparison with SCR control) in both fractions, but more significantly in the insoluble fraction. (B and C) Quantification of the ASO effects on total ubiquitin levels in the insoluble (B) and soluble (C) fraction of WT and TAR4/4 brain lysates. Group differences were found by ANOVA F_(3,12)_= 70.86, p<0.0001, post hoc analysis: ***, p<0.001 **, p<0.01, ***, p<0.001 (B) and ANOVA F_(3,54)_= 20.65, p<0.0001, post hoc analysis: ***, p<0.001 ***, p<0.001 (C). (D) Proteasome activity was assessed by measuring the hydrolysis of fluorogenic suc-LLVY-AMC (12.5 μM) by purified human proteasomes (10 nM) in the absence and presence of sarkosyl-soluble or -insoluble fractions from whole-cell lysates of TDP43-Q331K-overexpressing HEK293 cells. Time-dependent increase in activity of purified proteasomes (black circles) was completely inhibited by incubation with MG132 (black squares). Incubation of proteasomes with the sarkosyl soluble fraction of TDP-43 expressing HEK293 cells led to a modest reduction in proteasome activity (blue triangles) that was not statistically significant. Incubation of proteasomes with the sarkosyl insoluble fraction of TDP-43 expressing HEK293 cells led to a robust reduction in proteasome activity (red triangles) that was statistically significant. (E) Quantification of proteasome activity at the 60 minute time point reveals groups differences by ANOVA (F_(3,8)_=72.8, p<0.0001). The post hoc analysis shows that the proteasome activity in the sarkosyl-insoluble treated samples was statistically significantly reduced in comparison with proteasomes untreated with cell lysates (***, p<0.0001) or treated with sarkosyl-soluble lysates (**, p<0.001). Exact p values and confidence intervals are summarized in Supplementary Figure 7. (F) Affinity purified 26S proteasomes contain PSMD4, PSMA4, PSMB6 and RAD23A (by immunoblots) while 20S proteasome lack RAD23A. (G_i_) Whole cell lysate (WCL) from the brain of WT and TAR4/4 animals treated with SCR or RAD23A targeting ASO were assayed for proteasomal protease activity using the LLVY peptide. There is reduced protease activity in the TAR4/4 SCR ASO treated mice in comparison with the other experimental groups. Group differences were found by ANOVA F_(3,53)_= 30.13, p<0.0001, post hoc analysis: ***, p<0.001 *, P,0.05, **, p<0.01, ***, p<0.001. (G_ii_) Quantification of brain lysate protease activity using the RPPGFSAFK peptide. Group differences were found by ANOVA F_(3,44)_= 21.0, p<0.0001, post hoc analysis: ***, p<0.001 **, p<0.01. (G_iii_) Quantification of brain lysate protease activity using the KKVAPYPME peptide. Group differences were found by ANOVA F_(3,54)_= 20.65, p<0.0001, post hoc analysis: ***, p<0.001 *, P,0.05, **, p<0.01, ***, p<0.001. (H) Immunoblots for proteasome components (e.g., PSMA3, PSMC3 and PSMC5) in the soluble and insoluble fractions of WT and TAR4/4 brain lysates treated with SCR or RAD23A targeting ASO. In WT animals proteasome components are predominantly found in the soluble fraction and ASO treatments do not affect their abundance. In contrast, in the TAR4/4 mice, the abundance of the proteasome subunits in the insoluble fraction is increased (in comparison with WT). ASO targeting rad23 reduces (in comparison with SCR control) the partitioning of proteasome subunits into the insoluble fraction. (I) Quantification of the PSMA3 abundance in the insoluble fraction. Group differences were found by ANOVA F_(3,12)_= 32.92, p<0.0001, post hoc analysis: ***, p<0.001 **, p<0.01, ***, p<0.001. (J) Quantification of the PSMC3 abundance in the insoluble fraction. Group differences were found by ANOVA F_(3,12)_= 42.15, p<0.0001, post hoc analysis: ***, p<0.001 **, p<0.01, ***, p<0.001. (K) Quantification of the PSMC5 abundance in the insoluble fraction. Group differences were found by ANOVA F_(3,12)_= 19.65, p<0.0001, post hoc analysis: ***, p<0.001 **, p<0.01, ***, p<0.001. (L) Immunoblots of immunoprecipitated material from cells expressing epitope tagged A315T hTDP-43, RAD23A or ubiquitin. The results from IP with antibodies to the epitope tagged protein were compared with IP using a control IgG from the same species. Cell were treated with MG132 or vehicle (dimethyl sulfoxide - DMSO), or MG132. Input (10% of initial material) reveals the presence of the transgene and all the interrogated proteins. IP’ed TDP-43 (with anti-FLAG) pulls down TDP-43 but no RAD23A (probing with either anti-V5 or -RAD23A). MG132 leads to a high molecular weight smear in the anti-ubiquitin blot (using anti-HA) which likely represents ubiquitinated TDP-43. IP’ed RAD23A (with anti-V5) pulls down RAD23A (using anti-V5 or -RAD23A. With a long exposure there is a faint TDP-43 band in the (blotting with anti-FLAG or -TDP-43) in the MG132 treated cells. The molecular weight of this band corresponds to non-ubiquitinated TDP-43. IP’ed ubiquitin (with anti-HA) pulls down ubiquitin and TDP-43 but not RAD23A and MG132 increases the ubiquitin and TDP-43 signal.

To investigate this possibility, we monitored degradation kinetics of fluorescent LLVY-AMC proteasome substrate by affinity-purified proteasomes in the presence of soluble and insoluble fractions of cell lysates expressing hTDP-43. We used cell lysates from HEK293 cells engineered to inducibly-express hemaglutanin (HA)-tagged mutant TDP-43 (Q331K) as these cells develop sarkosyl-insoluble TDP-43. After induction of HA-mutTDP-43, sarkosyl-soluble and-insoluble fractions were generated, normalized to TDP-43 levels and added to proteasomes along with LLVY-AMC. MG-132-sensitive proteasomal activity was seen in this assay and addition of the sarkosyl-soluble fraction of mutTDP-43 expressing cells led to a modest (not statistically significant) decrease in proteasome activity (statistical analysis in Supplemental Figure 10). In contrast the sarkosyl-insoluble fraction of mutTDP-43 expressing cells inhibited proteasomal protease activity at all time points and was ∼50% reduced at the one hour time point (Figure 5D,E, statistical analysis in Supplemental Figure 10). Immunoblots of purified proteasomes reveals a robust signal for PSMD4, PSMA4, PSMB6 and interestingly RAD23A (Figure 5F). Next we assayed proteosomal protease activity of TAR4/4 brain lysates using three different fluorescent substrates; LLVY-AMC and two longer peptides (e.g., RPPGFSAFK and KKVAPYPME) that are cleaved by only 26S proteasomes and report “trypsin-like” and “caspase-like” activities respectively^28^. TAR4/4 brain lysate degradation of LLVY-AMC was inhibited by MG132 and KKVAPYPME by increasing salt concentrations (which disrupt the interaction between the 20S catalytic particle from the 19S regulatory particle) thus ensuring that these substrates report ubiquitin-dependent proteasomal activity (Supplemental Figure 11). TAR4/4 lysates displayed reduced levels of proteolysis of all three substrates (TAR4/4;RAD23A^SCR^ versus WT;RAD23A^SCR^ ∼30%, ∼20%, 20%, p<0.001, Figure 5G_i__-iii_). These observations support the view that the hTDP-43 evokes sarkosyl-insoluble cellular fraction that inhibits the 26S proteasome (Figure 5A-C).

Several investigations have previously shown that neurodegenerative disease-causing proteins can lead to inhibition of proteasome function^29–35^, including tau^36^, PrP^Sc30^ and the glycine-alanine dipeptide repeat protein^35^. It has been proposed that a physical interaction (either direct or indirect) of disease proteins with the proteasome is the basis of activity inhibition. Since RAD23A binds both ubiquitinated substrates and the proteasome directly, we wondered whether RAD23A tethers ubiquitinated aggregates to the proteasome in a manner that poisons proteasome function. This model predicts that the proteasome would be found in the insoluble fraction of TAR4/4 brain lysates in a RAD23A-dependent manner. When we probed lysates from the TAR4/4 mouse (± RAD23A KD) for components of the proteasome, we detect PSMA3 (20S subunit alpha 3), PSMC2 (19S subunit, ATPase 3) and PSMC5 (19S subunit, ATPase 5) and they are present in higher abundance in the sarkosyl-insoluble, but not soluble, fraction (Figure 5 H-K); ∼150%, 50% and 75% respectively in the TAR4/4;RAD23A^SCR^ versus WT;RAD23A^SCR^ animals (p<0.001 for all three proteins). Nuclear factor erythroid-derived 2-related factor 1 (Nrf1/NFE2L1) is the master regulator of proteasome subunit expression upon proteasome impairment^37^ and its abundance is ∼ 10x higher in the brain TAR4/4;RAD23A^SCR^ versus WT;RAD23A^SCR^ animals (p<0.001). Knockdown of *RAD23A*, reduced the partitioning of the three proteasome proteins into the insoluble fraction by 45%, 20% and 45% respectively (p<0.01 all three proteins, Figure 5I-K) and reduces Nrf1 levels ∼ 50% (p<0.001, Supplemental Figure 12). Thus proteasome subunits incorporate into sarkosyl-insoluble aggregates evoked by hTDP-43 in a RAD23A-dependent manner.

The reduced sequestration of proteasome components in the insoluble fraction in the RAD23A KD brains might support the production of more functional proteasomes in the soluble fraction of cell lysates. In favor of this view while the TAR4/4 lysates displayed reduced levels of proteolysis of the three fluorogenic reporters, KD of RAD23A restored proteolytic activity ∼25%, ∼35% and ∼15% (TAR4/4;RAD23A^KD^ versus TAR4/4;RAD23A^SCR^, p<0.001, p<0.01 and p<0.001, respectively) (Figure 5G_i__–iii_). These observations suggest that the burden of non-degraded ubiquitinated proteins is high in the TAR4/4 brain for two, potentially distinct, reasons: 1) the sarkosyl-insoluble fraction of TDP-43 overexpressing cells directly inhibits the proteolytic activity of the proteasome and 2) RAD23A promotes the partitioning of proteasomes into insoluble aggregates where they display impaired proteolytic function.

Since hTDP-43 has a predilection to misfold and aggregate, it might initiate a cascade of protein aggregation → proteasome sequestration → reduced proteasome function → more protein aggregation. The role of RAD23A in this vicious cycle might be to link TDP-43 with proteasomes. If this is true, a physical association of TDP-43 and RAD23A should be demonstrable. To examine this possibility, we heterologously expressed epitope tagged proteins (e.g., HA-Ubiquitin (HA-Ub), FLAG-TDP-43 and V5-RAD23A), treated with proteasome inhibitor MG132 or vehicle and immunoprecipitated (IPed) with anti-epitope tag or irrelevant IgG beads (Figure 5L). IP of TDP-43 (with anti-FLAG) pulled down TDP-43 (blotting with anti-FLAG or -TDP-43) and a high molecular weight smear of ubiquitin (blotting with anti-HA) only after MG132(+) (likely polyubiquitinated TDP-43). There is no signal in the anti-V5 blots indicating that RAD23A is not found in the TDP-43 pull down. In the reciprocal experiment, IP of RAD23A (using anti-V5) revealed RAD23A (blotting with anti-V5 or -RAD23A) and in long exposure imaging there is a faint band for TDP-43 in the MG132(+) lysates (blotting for FLAG or TDP-43). Thus a minor fraction of overexpressed TDP-43 physically associates with RAD23A upon proteasome inhibition. We wondered if enriching for ubiquitinated species would reveal a more robust TDP343-RAD23A association. We IPed ubiquitin (with anti-HA) and detected a smear of high molecular weight ubiquitinated proteins when probing for ubiquitin. We see a band consistent with monoubiquitinated TDP-43 (when probing with anti-FLAG or-TDP-43) but only in the MG132(+) lysates. Probing for RAD23A reveals a faint band in the MG132 lysates consistent with RAD23A binding some ubiquitinated proteins (as expected) or ubiquitinated RAD23A itself. Together the data does not support a robust physical link of RAD23A to ubiquitinated TDP-43. This raises the question: how does KD of RAD23A reduce the association of proteosomes with TDP-43-evoked aggregates? We posit that KD of RAD23A leads to a qualitative alteration in the composition of the sarkosyl-insoluble fraction of cell lysates evoked by TDP-43. While protein aggregates supervised by RAD23A have a propensity to sequester proteasomes this apparently occurs absent a direct physical interaction between RAD23A and TDP-43.

Beyond the proteasome, are other cell biological processes affected by RAD23A-dependent changes in the composition of the insoluble fraction of cells? Prior mass spectrometry interrogation of the insoluble proteome in NDD have shown enrichment of protein homeostasis machinery (i.e., proteasome subunits^33,38–40^, components of aggresomes^33,38^ and autophagy^33,41^) and nuclear complexes/splicing^33,38,39,41^. Here, we used stable heavy isotope (^15^N)-based quantitative liquid chromatography-tandem mass spectrometry (LC-MS/MS) based proteomics to compare the protein composition of the insoluble fraction of TAR4/4;RAD23A^KD^ versus TAR4/4;RAD23A^SCR^ brain lysates(Figure 6A). We quantified 6,343 protein IDs and the majority were not elevated in one experimental group. 304 protein IDs were differentially modulated; 146 in the SCR treated brains and 158 in the RAD23A KD brains (at ≥1.5 fold change and p-value <0.05). Representative MS1 spectra are shown in Figure 6B. A t-distributed stochastic neighbor embedding (tSNE) plot shows that the two experimental groups are distinct indicating that differentially enriched proteins are consistent among experimental groups (Figure 6C). A volcano plot illustrates differentially insoluble proteins as a function of abundance and statistical significance (Figure 6D). Consistent with our immunoblot data, TDP-43 is more enriched in the SCR ASO treated versus RAD23A ASO treated samples. Similarly, multiple examples of proteasome components and interacting proteins (i.e. PSMD2, HSPA2, SNAP25, DNAJ, ENO1, etc.)^42,43^ are enriched in the SCR ASO treated versus RAD23A treated samples consistent with RAD23A-dependent sequestration of proteasomes as described above (Figure 5G-I). More broadly, these results are evidence that RAD23A governs molecular composition of the insoluble proteome evoked by expression of TDP-43.

**Figure 6.**
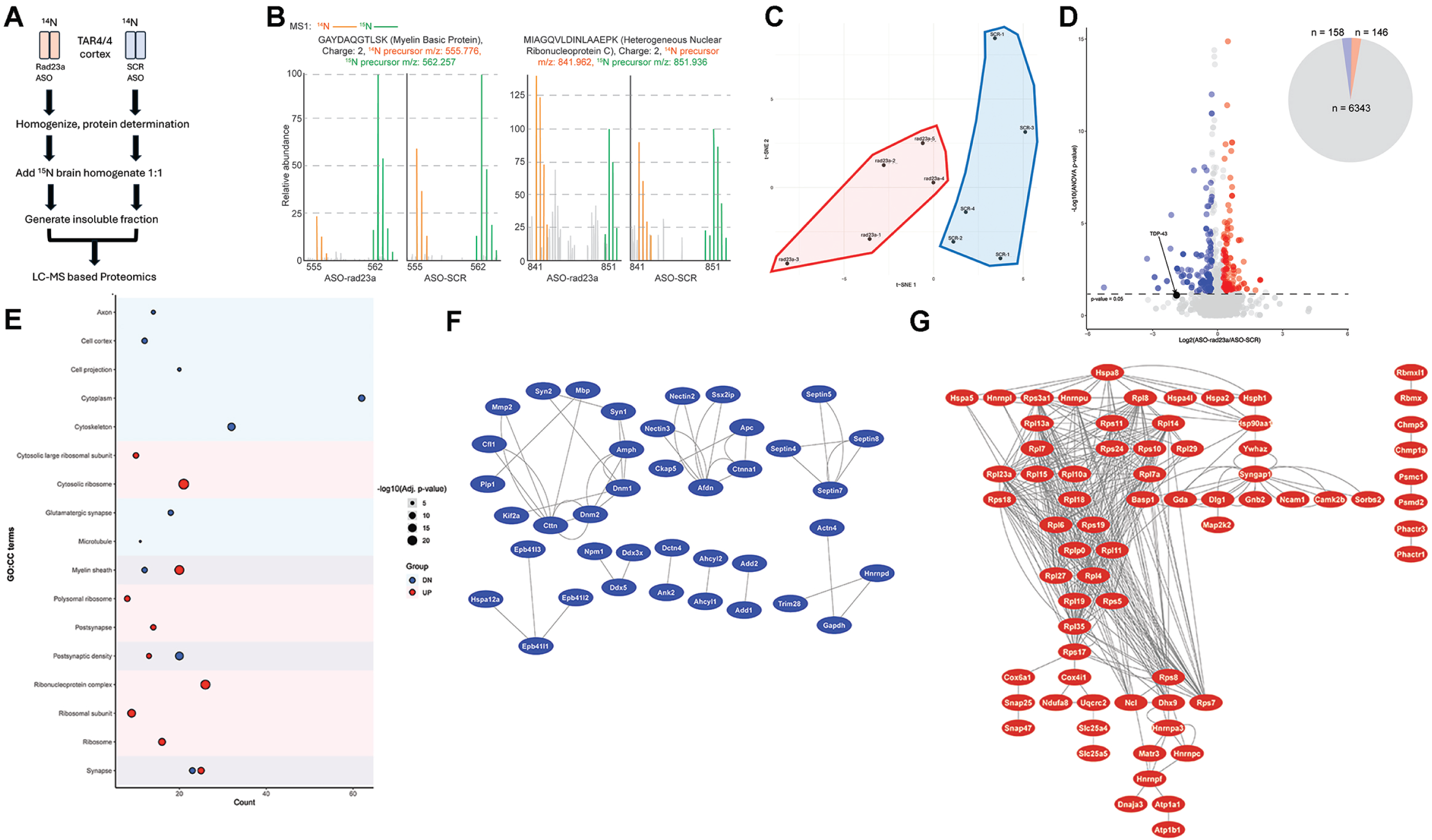
RAD23A controls the composition of the insoluble protein fraction evoked by hTDP-43 expression in mice. (A) Cartoon of experiment. TAR4/4 animals on standard show (i.e., ^14^N unlabeled proteins) received the ASO targeting RAD23A or the scrambled sequence ASO. Motor cortex was homogenized and protein concentration determined. In parallel motor cortex the ^15^N fully-labeled mouse was homogenized and protein content determined. A 1:1 mix of ^14^N with ^15^N homogenates was created for each experimental sample and then soluble/insoluble fractions generated by centrifugation. The insoluble fraction was then recovered and subjected to the LC-MS analysis. (B) Annotated representative raw MS1 spectra across the specified m/z ranges display distinct peptides from myelin basic protein and heterogeneous nuclear ribonucleoprotein C. The spectra highlight ^15^N peptide peaks in green, ^14^N peptide peaks in orange, and other peaks in gray. (C) Biological replicates cluster according to treatments in the tSNE (t-distributed stochastic neighbor embedding) plot. (D) Volcano plot comparing the insoluble protein fraction components of mice receiving RAD23A knockdown versus scrambled sequence. Peptide-based ANOVA test. Pie chart illustrating the differences between treatments. (E) GO cellular component (CC) terms of proteins significantly enriched in either experimental groups. X-Axis, GO terms, Y-axis, counts (greater values indicate a larger number of proteins within the GO term). Blue circles – significantly reduced enrichment in RAD23A ASO treated versus SCR ASO treated, Pink circles – significantly enhanced enrichment in RAD23A ASO treated versus SCR ASO treated. Circle size reflects -Log10 adjusted p-values. (F) Protein-protein interaction (PPI) network of proteins with significantly reduced enrichment in RAD23A ASO treated samples compared to SCR ASO treated samples. (G) PPI network of proteins with significantly increased enrichment in RAD23A ASO treated samples compared to SCR ASO treated samples.

A GSEA of the proteins differentially enriched in the insoluble proteome ±RAD23A was performed (Figure 6E). A prominent set of differentially enriched proteins that were higher TAR4/4;RAD23A^SCR^ versus TAR4/4;RAD23A^KD^ were cytoskeletal and motors (i.e. septins 4, 5, 7, 8, actin, cortactin, dynamin 1, 2, dynactin 4, Kinesin 2a, etc.). These proteins are part of functional protein association networks as revealed in GeneMania protein-protein-interaction (PPI) network analysis (Figure 6F). This observation is consistent with the known cytoskeletal abnormalities as cardinal features of NDDs and likely of pathophysiological significance. The prominent set of differentially enriched proteins in the TAR4/4;RAD23A^KD^ versus TAR4/4;RAD23A^SCR^ are ribosomal (i.e., Rpl 1, 3-8, 10-15 etc.) and the PPI network analysis reveals dense physical interactions (Figure 6G). If this enrichment coincided with a general reduction in protein synthesis this might be beneficial by better aligning nascent protein synthesis with protein folding capacity. RAD23A influences the composition of the insoluble proteome of evoked by hTDP-43 and this likely contributes to the beneficial actions of reducing RAD23A levels.

## Discussion

Here we show that reducing the abundance of RAD23A blunts the toxicity of human TDP-43 overexpression *in vivo*. The beneficial effects on lifespan, motor behavior, histological abnormalities and distortion of the transcriptome are likely derived from restoration of the nuclear TDP-43 activity, increased proteasome function and remodeling of the protein composition of the insoluble proteome. At least part of the beneficial effects appears to be driven by reduced sequestration of specific proteins that are components of functional networks that go awry in NDD.

Aggregates of misfolded TDP-43 can be toxic agents in a variety of ways. TDP-43 aggregates can form a nidus that incorporates other cellular proteins into an insoluble fraction causing cellular depletion by sequestration (i.e., nuclear pore complex and nucleocytoplasmic transport machinery^39^ and proteins involved in RNA metabolism and proteasome subunits^33^). Aggregates of misfolded proteins can also disrupt protein homeostasis on a more global level by monopolizing chaperones, for example, and interfering with the normal function of a sub-proteome of metastable proteins^44,45^. Prolonged engagement of the proteasome with misfolded proteins (perhaps due to slow unfolding and processive degradation) can also lead to neglect of other ubiquitinated substrates^31^ and their accumulation. Thus by virtue of the propensity of TDP-43 to misfold and aggregate it can initiate broad adverse effects on the integrity of the proteome. Seminal work by the Polymenidou group has shown that TDP-43 aggregates display different sizes, morphologies, densities, co-aggregating proteins and neurotoxicities^46^. Owing to low complexity domains, portions of TDP-43 can adopt a variety of structures with low energy barriers between conformations. It is possible that diversity of TDP-43 conformers is a driver of the compositional heterogeneity of aggregates.

The data presented here points to RAD23A as another determinant of aggregate composition but by a mechanism that does not depend on a physical interaction of TDP-43 and RAD23A. How might this work? Protein unfolding is required for proteolysis by the proteasome and specific physical features of the substrate govern this process. Exquisite studies in yeast reveal that well-folded, ubiquitinated substrates with a flexible segment of sufficient length and orientation can be unfolded by the 19S regulatory particle of the proteasome. If, however, the flexible segment lacks favorable features, unfolding is performed by cdc-48 (VCP in vertebrates) prior to engagement with the proteasome. RAD23 and Ubx5 (Ubxn7 in vertebrates) are the key cofactors directing substrates to the 19S particle or cdc-48/VCP, respectively. RAD23 inhibits the degradation of proteins that require unfolding by cdc48/VCP – they adhere to the proteasome in a non-productive orientation. In the absence of cdc48 to unfold a substrate, >1,000 proteins adhere to the proteasome in a RAD23-dependent manner. At least 75% of this pool of proteins can be dissociated from the proteasome by overexpression of Ubx5. Thus a very substantial portion of the proteome shuttles between the proteasome and cdc48/VCP as a function of cofactor proteins like RAD23 and Ubx5. The balance of this shuttling governs the unfolding/degradation versus accumulation of ubiquitinated substrates. The beneficial effects of RAD23A KD that we observe may derive from an improved balance of substrate assignment to VCP prior to proteasomal degradation. Future work will be required to test this idea.

TDP-43 pathology is commonly seen in a variety of NDD and is likely to be a key pathophysiological driver. Since TDP-43 is an essential protein for cellular function, therapeutically targeting its pathological states is challenging. We provide evidence here that RAD23A controls TDP-43 toxicity by sculpting UPS function and creating a more favorable proteomic landscape within neurons. If ASOs targeting human RAD23A could be developed, there is a clear a path for appraising their utility in human NDD.

## Methods and Materials

### Mouse crosses

All procedures related to the animal study performed here were approved by the Northwestern University Feinberg School of Medicine Institutional Animal Use and Care committee. All experiments were conducted in accordance with USDA Animal Welfare Act and PHS Policy on Humane Care and Use of Laboratory Animals. Up to 5 animals per cage were housed together under a 12 hr light/dark cycle. TAR4 mice (*TDP-43^Tg/+^*) were purchased from Jackson Laboratory (Stock #012836, B6;SJL-Tg(Thy1-TARDBP)4Singh/J). They were maintained on a B5/SJL hybrid background breeding with B6/SJLF1/J (JAX Stock #100012). Genotyping was performed as published^26^ and as described in the JAX protocol. RAD23A knock out mice were generated in the R.G.K lab, described in detail below. They were back crossed in the C57Bl6 background > 8 generations. The line was maintained by mating *RAD23A*^+/-^ with non-littermate C57Bl6 mice. Experimental animals for the ASO studies were generated by intercrossing non-sibling *TDP-43^Tg/+^* with genotype establish *post hoc*. For the studies of genetic manipulation of RAD23A levels, the *Rad23^+/-^*mice were bred to *TDP-43^Tg/+^* to generate mice with genotypes *RAD23A^+/+^; TDP-43^Tg/Tg^, RAD23A^+/-^;TDP-43^Tg/Tg^* and *RAD23A^-/-^;TDP-43^Tg/Tg^.* We predicted an effect size, if it existed, would mimic that seen by Becker et al.^47^ and therefore instead of a power analysis with simply estimated that ∼20 animals per genotype would be needed in the lifespan, gait interrogations and the analysis of other phenotypes. The ASO survival study was performed on two independently generated cohorts of animals created more than one year apart.

### Generation of the RAD23A knock out allele

RAD23A KO mice (genotype as *RAD23A*^-/-^) were generated using CRISPR/CAS9 targeting nucleotides in NC_000074.7, exon 7 to create a stop codon. The nucleotides flanking the edit were “cagcaggggagaaccccctg(GA/C)gttcctgcgggaccagcctc” with GA edited to C. Genotyping was performed with primers 5’-CTGAACATGGTTCTGTCCAG-3’ and 5’-CCCCTAAGGAACTGGGAAG-3’ to amplify a 339 bp PCR product, which was then digested with BstN1 to release a 268bp fragment specific for RAD23A KO. Western blot was also performed to confirm the loss of RAD23A at protein level in transgenic mice.

### ASOs validation and administration

ASOs that selectively target either mouse Rad23a or Rad23b were designed, synthesized and screened by Ionis Pharmaceuticals. Synthesis and purification of all chemically modified oligonucleotides were performed as previously described^48^. The ASOs are 20 nucleotides in length, in which the central gap segment has ten 2′-deoxyribonucleotides that are flanked on either side by five 2’MOE modified nucleotides. Internucleotide linkages are phosphorothioate interspersed with phosphodiester, and all cytosine residues are 5′-methylcytosines.The following ASOs with high efficiency and well tolerance were selected for this study—RAD23A, CCTTGCTTATTATACCAAAC; and RAD23B, ACATATTATTTCAGACGCAC. One ASO with scramble sequence (SCR), CCTATAGGACTATCCAGGAA, was included as negative control.

Postnatal day zero (P0) neonates were used for ICV injection with 15-60 μg ASOs in 3 μl sterile phosphate buffered saline PBS as described. The most effective and well tolerated dose of each ASO was determined empirically. In brief, pups were transferred to ice-cold metal plate to induce hypothermia anesthesia and the head was cleaned with a 70% ethanol cotton swab prior to injection. The injection site was identified as 2/5 of the distance from the lambda suture to each eye. A 32 Gauge Hamilton syringe was held perpendicular to the surface of the skull and the needle was penetrated the skin and skull to a depth of ∼3 mm. ASOs were slowly delivered into the lateral ventricle by depressing the plunger. The injection was conducted on both sides with 1.5 μl administrated into each hemisphere. Before returning to the nest, the pup was transferred onto a warming pad until body temperature and color normalized and the animal was moving freely. Half pups of the litter were randomly chosen for *RAD23A* or *RAD23B* ASO injection, while the other half received the scramble sequence ASO.

To obviate bias, ICV injections were performed with the operator blind to genotype and whether the ASO was SCR sequence or designed to target RAD23A or RAD23B. Three microliters of ASO were loaded in a coded manner into 8-strip PCR tubes by experimentalist 1. Experimentalist 2 then took the 8-strip PCR tube and loaded the syringe for injection of new born mice. Each mouse received a tattoo to allow for linking of animal to ASO injection and subsequent genotyping (Aramis Laboratory animal microtattoo system, Ketchum # 623). These animals were then used for behavioral phenotypes and lifespan studies. Upon euthanasia a tail snip was obtained and PCR genotyping performed. The coordination of genotype with ASO treatment was not revealed until data was analyzed.

The tissue samples for RNAseq analysis were obtained by experimentalist 1 and analyzed by a second experimentalist blind to genotype and ASO. Once the RNAseq data was acquired and analyzed, the animal groups were revealed to experimentalist 2.

### Care of the TAR4/4 (TDP-43^Tg/Tg^) mice

Breeding pairs were feed on high-fat breeder chow (11.4% fat; Inotiv #7904). If TAR4/4 pups reached the humane endpoint before P21, they were removed euthanasia. If not, pups were weaned separately into new cages at P21 to avoid injury of the TAR4/4 mice by their littermates or parents. To ensure adequate nutrition to support the survival of compromised TAR4/4, wet food DietGel® 76A (w/Wheat Protein) from ClearH2O was provided on the cage floor in addition to the moisturized chow and bottled water. Food was changed every day to keep fresh. This ensured that all mice were able of healthful survival before reaching the humane endpoint.

### Phenotype scoring and humane euthanasia end point determination

Behavior and lifespan study were performed as described^47^. Behavioral features we studied include hind limb clasping, gait and kyphosis. The assessment of animal behavior was performed blind to the animal genotype or treatment (ASO SCR versus targeting *RAD23A* or *RAD23B*). Behavioral assessments were made on animals placed on a warm, textured surface not in the cage setting. Gait impairment scoring system measures motor dysfunction and as demonstrated by Becker et al. suitable for study of young mice. Animals that walk normally without visible impairment were assigned a score of 0; animals walking with a slight tremor or limp were assigned a score of 1; animals walking with severe tremor, severe limp, lowered pelvis, or feet splayed away from the midline (“duck feet”) were assigned a score of 2; animals displaying troubled forward movement, minimal joint movement, and poor utilization to generate forward movement and stayed upright with difficulty (or dragged their abdomen on the examination surface) were assigned a score of 3; animals that were unable to maintain an upright posture and unable to right within 30 sec on 3 trials was assigned a score of 4. Behavioral assessments were typically made every other day from 14 of life. Once a score of three was reached, the animals were assessed daily. Animals that reached the score of 4 had reached the humane pre-specified end point for the lifespan study and were euthanized.

We used the kyphosis and tremor scoring system developed by Becker et al.^47^ For kyphosis, animal without visible dorsal curvature of the spine and who are able to easily straighten its spine while walking were assigned a score of 0; Score 1 means a animals with mild kyphosis but still able to straighten its spine while walking were assigned score of 1; animals with a persistent but mild kyphosis and unable to completely straighten its spine most of the time were assigned a score of 2; animals with pronounced kyphosis at all times were assigned a score of 3. For tremor, animals with no detectable shaking were assigned a score of 0; Score 1, animals with very mild shaking or occasional shaking were assigned a score of 1; animals with severe but not continuous shaking were assigned a score of 2; shaking severely but not continuously; animals with continuous severe shaking were assigned a score of 3. Hindlimb clasping was assessed by the gently grasping the mouse near the tail base and lifted to observe the hindlimb position for 10 sec. Score 0 means both hindlimbs are consistently splayed out; Score 1 means one hindlimb is retracted toward the abdomen for half time of suspension or more while another hindlimb still splaying outward all the time; Score 2 means both hindlimbs are loosely retracted for half the time of suspension or more; Score 3 means both hindlimbs are closely or tightly retracted for half the time of suspension or more.

### Mouse tissue collection

Deeply anesthetized mice were exsanguinated by cardiac laceration and the central nervous system (brain and spinal cord) were rapidly dissected out of the corpse. For biochemical studies, the brain was regionally subdivided (e.g., cortex, hippocampus and remaining regions), frozen in dry ice and stored wrapped in tin foil with a coded designation. For transcriptomic and proteomic studies one half of the brain was subjected western blot to confirm efficacy of ASO treatment. For immunohistological studies, the brain was fixed by immersion in freshly prepared 4% paraformaldehyde in 0.1 M pH=7.4 phosphate buffer overnight and then washed extensively in phosphate buffered saline (0.1 M phosphate pH=7.4 buffer + 0.9% sodium chloride) prior to further processing.

### Total RNA extraction

The motor cortices of WT, TAR4/4 and Rad23 ASOs treated mice were dissected from left hemispheres of mice. The RNASeq library preparation and sequencing were carried out by the Genewiz facility(https://www.genewiz.com/en-GB/). Total RNA was extracted (Qiagen cat. No. 1023539), quantified (Qubit 4.0 Fluorometer (Invitrogen, Carlsbad, CA) and RNA integrity was checked with TapeStation 4200. RNA sequencing libraries were prepared using the NEBNext Ultra II RNA Library Prep Kit (NEB, Ipswich, MA, USA). Briefly, mRNA was enriched with Oligo(dT) beads and fragmented according to the manufacturer’s instructions before first-strand and second-strand cDNA synthesis. cDNA fragments were end-repaired, adenylated at 3’ends, ligated with universal adapters, followed by indexing and enrichment by limited-cycle PCR. Libraries were validated using the NGS Kit on an Agilent 5300 Fragment Analyzer (Agilent Technologies, Palo Alto, CA, USA), quantified (Qubit 4.0 Fluorometer), multiplexed and loaded on flow cells for sequencing (Illumina NovaSeq 6000) using a 2x150 paired-end configuration v1.5. Image analysis and base calling were conducted by the NovaSeq Control Software v1.7. Raw sequence data (.bcl files) was converted into fastq files and de-multiplexed (Illumina bcl2fastq program version 2.20). One mismatch was allowed for index sequence identification.

### RNA-Seq data analysis

Low-quality reads (Phred<20), and adapters were removed using Trim-galore (V0.6.7) with its standard parameters (https://github.com/FelixKrueger/TrimGalore.com/fenderglass/Flye). Afterwards, reads were aligned to the GRCm38/mm10 build using RNA STAR Gapped-read mapper (V-2.7.11a+galaxy1 or V-2.7.8a+galaxy1)^49^. The alignment was carried out using standard parameters without transcript guidance(GTF) using the galaxy platform^50^. To quantify expressed genes, we counted mapped reads on exons (GRCm38.99.gtf) using featureCounts(V 2.0.1+galaxy2)^51^. The differentially expressed genes between the two conditions were measured using DSeq2^52^. Lowly expressed genes (reads < 10) were filtered, and samples were normalised using the median of ratio method. A parametric fit was used to reduce the influence of false-positive outliers on fold change and p-value. All genes with an adjusted p-value (FDR) of less than 0.05 were considered to be differentially expressed. For the downstream analysis, read counts of all the samples was normalized by DESeq2 using the same median ratio method as discussed earlier. The rlog of normalised read counts was used to perform principal component analysis (PCA), gene set enrichment analysis (GSEA) and hierarchical clustering. GSEA was performed using GSEA_4.3.2, M2.all.v2023.2mm curated genes and normalised read counts^53^. All genes with normalised read counts > 1 were included in the analysis. PCA, Volcano plot and heatmaps were generated using R-language (https://www.r-project.org/help.html.

### Triton X-solubility

Protein was extracted from frozen tissues as indicated in the main text. Tissues were snap-frozen and were weighted, which was recorded as initial weight. All subsequent volumes were based on this initial weight. For general western blotting, tissues were homogenized by 50 strokes in Kontes with 10 Vol RIPA buffer supplemented with 1 X protease inhibitor (Sigma) and 1 X phosphatase inhibitor (PhosphoStop, Roche). The homogenates were incubated on ice for 10 min and then centrifugated at 18,000 g for 20 min at 4°C. The supernatant was saved as Triton X-soluble fraction and the pellet was washed with the extraction buffer and centrifugated again as above. The second round pellet was dissolved in 2 Vol urea buffer (30 mM Tris, pH 8.5, 4% CHAPS, 7 m urea, 2 m thiourea) as Triton X-insoluble fraction.

### Immunoblotting

Approximately 30 μg protein was heated at 85°C for 5 min with 1 X SDS loading buffer. The samples were loaded onto 10% or 12.5% SDS-PAGE gel depending on the size of target proteins. After it, the protein was transferred overnight onto nitrocellulose membrane. The membranes were then blocked, probed overnight at 4°C with the primary antibodies, washed 3 times in TBST, and incubated with appropriate secondary antibodies conjugated with Alex Fluor-680 or - 800. All the antibodies used in this study was listed in Extended Table 2. Imaging was realized with Li-Cor Odyssey scanner and protein quantification was performed in Image Studio Lite Version 5.2.

### Cytoplasmic and nuclear fractionation

Nucleocytoplasmic fractionation was conducted as described^26,47,54,55^. Briefly, tissue was homogenized with Kontes in 10 vol/wt hypotonic buffer (10 mM Hepes, 10 mm NaCl, 1 mm KH_2_PO_4_, 5 mm NaHCO_3_, 5 mm EDTA, 1 mM CaCl_2_, 0.5 mm MgCl_2_ with 1 X protease inhibitor and 1 X phosphatase inhibitor). After homogenization, 0.5 vol/wt 2.5 m sucrose was added into the lysate before a second round homogenization. The homogenate was centrifuged at 6,300 g for 10 min. The supernatant was saved as the cytoplasmic fraction. The pellet was washed three times in pre-cold TSE buffer (10 mM Tris, pH 7.5, 300 mm sucrose, 1 mm EDTA, 0.1% NP-40) by resuspending and then centrifuging at 4,000 × g for 5 min. The pellet was dissolved in 2% SDS buffer and was saved as nuclear fraction.

### HA-mutTDP43 Q331K Flp-In T-Rex cell line

HA tagged human Q331K mutant TDP-43 was cloned in the pcDNA5/FRT/TO plasmid (Invitrogen™ V652020) and sequence verified. HEK293 Flp-In T-REx cells (ThermoFisher R78007) were cultured in DMEM (Gibco™ 11965092), 10% Tetracycline free FBS (Gibco™ A4736201), 1x penicillin-streptomycin (Gibco™ 15140122), 1X GlutaMAX™ (Gibco™ 35050061), 15 ug/mL Blasticidin S (Gibco™ A1113903) and 100 ug/mL zeocin (Gibco™ R25001). To create^56,57^ a stable inducible cell line, 3x10^5^ cells were plated per well of a 6-well plate and transfected when 70% confluent 48 hours later with 125 ng of pcDNA5/FRT/TO–HA-Q331K TDP-43, 1.125 ug pOG44 (Invitrogen™ V600520), and 6.25 uL Lipofectamine 2000 (Invitrogen™ 11668500) per well. After 48 hours, cells were re-plated in media without zeocin and with 150 ug/mL hygromycin B (Gibco™ 10687010) for selection. Cells were selected for 3 weeks before re-plating for expansion and screening for successful integration via immunoblotting.

### SarkoSpin

SarkoSpin was performed as described before^46,58^. Tissues were homogenized in homogenization solubilization (HS) buffer (10 mm Tris, pH 7.5, 150 mm NaCl, 0.1 mM EDTA, 1 mm DTT, 1X protease inhibitor (Roche) and Phosphatase inhibitor (Roche)). Samples were then diluted in 1 Vol of HS buffer with 4% Sarkosyl containing 2U/μl benzonase and 4 mm MgCl2. After incubating at 37°C for 45 min under constant shaking at 600 rpm, brain homogenates were further diluted with 0.5 Vol HS buffer containing 0.5% Sarkosyl. Samples were then centrifugated at 21,000 g for 30 min to separate Sarkosyl-soluble and -insoluble fractions. Supernatants were collected in a new tube and saved as Sarkosyl-soluble fraction. Pellets were washed by resuspeneding in HS buffer with 2% Sarkosyl and spinning down again at 21,000 g for 30 min to obtain Sarkosyl-insoluble fraction, which were dissolved in 2% SDS for western blot analysis.

### Immunohistochemistry

Immunohistochemistry was performed according to standard protocol^59^. Paraffin-embedded 6 µm-thick sections were deparaffinized in xylene and rehydrated in graded alcohol concentrations. Antigen retrieval was performed in 1% citrate buffer heat retrieval solution, pH 6.0 (Antigen Decloaker, Biocare Medical) using an electric pressure cooker at 121°C for 20 minutes. Endogenous peroxidase activity was blocked with 3% hydrogen peroxide for 20 minutes. To reduce nonspecific signals, sections were blocked with 1% Bovine Serum Albumin (BSA) in 1x Phosphate-buffered saline (PBS). Sections were incubated with primary antibodies diluted in blocking buffer overnight at 4°C in a humidified chamber. Sections were incubated with peroxidase-conjugated secondary antibody (Jackson ImmunoResearch) diluted in a blocking buffer for 2 hours at room temperature. 3,3′-Diaminobenzidine (DAB) solution was prepared by dissolving a DAB tablet (Sigma-Aldrich D4293) in 5 ml of deionized water containing 0.01% hydrogen peroxide. Sections were finally treated simultaneously with the DAB solution for 2-3 minutes at room temperature. The following antibodies were used: 1:500 rabbit NeuN (Abcam, ab104225), 1:250 mouse TARDBP (Abnova, H00023435-M01), 1:200 mouse Ubiquitin (Enzo, BML-PW0930), 1:200 rabbit Glial Fibrillary Acidic Protein (Dako, Z0334), 1:200 rabbit Iba1 (Wako, 019-19741)

### RNA extraction and RT-qPCR

Motor cortex tissues were dissected in RNase free conditions from freshly culled mice and flash frozen for gene transcription analysis^60^ here from different-treated mice. RT-qPCR was conducted as reported before^61^. Briefly, total RNA was extracted with the RNeasy® Plus Mini Kit (Qiagen #74134) following the manufacturer’s instruction. RNA integrity and quantity was determined by NanoDrop. Only those with an A260/A280 ratio above 1.95 were used for the following reverse transcription. The RNA was converted to cDNA using qScript cDNA SuperMix (Quantbio #95048-100). Real time qPCR was performed on StepOnePlus Real-Time PCR System (Applied Biosystems™) with Power SYBR™ Green PCR Master Mix (Thermofisher #4368706) and primers as listed in Extended Table 3. The amplification conditions comprised 50°C for 2m, 95°C for 10 min, followed by 40 cycles of “95°C for 10s, 60°C for 1 min”. Each reaction was performed in triplicate. GAPDH and β-actin were included as housekeeping genes and their average Ct values were used for normalization. The relative transcript levels were calculated by the method of ΔΔCt.

### Proteasome activity assay

For measuring proteasome activity^28^, brain cortex tissues were homogenized in proteasome lysis buffer (40 mM Tris, pH 7.5, 50 mM NaCl, 2 mM β-mercaptoethanol, 2 mM ATP, 5 mM MgCl_2_, 10% glycerol) to whole cell extracts. The lysates were briefly sonicated and then centrifugated at 21,000 g for 20 min at 4°C. Protein concentration of the supernatants containing cellular proteasomes were measured with BCA assay and was adjusted to 5 mg/ml for hydrolysis assay as follows. Whole proteasome activity was assessed with various fluorogenic substrates (all from Enzo Life Sciences), in reaction buffer (50 μM Tris-HCl, pH 7.5, 1 mg/ml BSA, 1 mM ATP and 1 mM DTT) at 37°C for 30 min, including 50 μM Suc-LLVY-Amc 20 μM Mca-RPPGFSAFK(Dnp) and 20 μM Mca-KKVAPYPME-Dap(Dnp)–NH2. Fluorescence was measured using a BioTek Synergy II plate reader with the excitation and emission (Ex/Em) filters at 360/460 nm for Amc and 320/405 nm for Mca

To evaluate the effect of insoluble TDP-43 on proteasome activity, a suc-LLVY-AMC hydrolysis assay was conducted for indicated time periods using 10 nM affinity-purified proteasomes^62^ with sarkosyl-soluble or -insoluble fractions from doxycycline (Dox)-inducible HEK293 cells. Cells were pre-treated with 2.5 μg/mL Dox for 24 h to induce TDP43-Q331K overexpression, and a modified SarkoSpin method was used to separate soluble and insoluble fractions of whole-cell lysates. The volumes of these fractions were adjusted so that the band intensities were equal in immunoblotting. Protein concentrations were determined using the Bradford method for soluble samples and gel-based TDP43 immunoblotting for insoluble samples. Similar experiments were performed using the inducible HEK293 cells after RAD23A was knocked-down by siRNA.

### MS sample preparation

The anterior half of the cerebrum that contains the motor cortex wsa homogenized in 500 uL of fresh, ice cold 0.32 M sucrose, 10 mM HEPES + protease inhibitors using Precellys beads (4,000 rpm x 15 seconds, one cycle). BCA protein determination was performed and 200ug of homogenate was combined 1:1 with 200 ug of ^15^N brain homogenate that was processed in parallel^63^ and the total column brought to 400 uL with homogenization solution. Forty uL of 10% SDS was added to each sample and equilibrated on a rotator at 4o C. for 30 minutes prior to centrifugation at 100,000*g* for 60 minutes. The supernatant was dischared and the protein pellets were resuspended in 8 M urea (ThermoFisher Scientific, Cat # 29700) prepared in 100 mM ammonium bicarbonate solution (Fluka, Cat # 09830) and processed with ProteaseMAX (Promega, Cat # V2072) according to the manufacturer’s protocol. The samples were reduced with 5 mM Tris(2-carboxyethyl)phosphine (TCEP, Sigma-Aldrich, Cat # C4706; vortexed for 1 hour at RT), alkylated in the dark with 10 mM iodoacetamide (IAA, Sigma-Aldrich, Cat # I1149; 20 min at RT), diluted with 100 mM ABC, and quenched with 25 mM TCEP. Samples were diluted with 100 mM ammonium bicarbonate solution, and digested with Trypsin (1:50, Promega, Cat # V5280) for overnight incubation at 37°C with intensive agitation. The next day, reaction was quenched by adding 1% trifluoroacetic acid (TFA, Fisher Scientific, O4902-100). The samples were desalted using Peptide Desalting Spin Columns (Thermo Fisher Scientific, Cat # 89882). All samples were vacuum centrifuged to dry.

### Tandem Mass spectrometry

Three micrograms of each fraction or sample were auto-sampler loaded with an UltiMate 3000 HPLC pump onto a vented Acclaim Pepmap 100, 75 m x 2 cm, nanoViper trap column coupled to a nanoViper analytical column (Thermo Fisher Scientific, Cat#: 164570, 3 µm, 100 Å, C18, 0.075 mm, 500 mm) with stainless steel emitter tip assembled on the Nanospray Flex Ion Source with a spray voltage of 2000 V. An Orbitrap Fusion (Thermo Fisher Scientific) was used to acquire all the MS spectral data. Buffer A contained 94.785% H_2_O with 5% ACN and 0.125% FA, and buffer B contained 99.875% ACN with 0.125% FA. The chromatographic run was for 2 hours in total with the following profile: 2–8% for 6, 8–24% for 64, 24–36% for 20, 36–55% for 10, 55–95% for 10, 95% for 10 mins.

We used CID-MS^2^ method for this experiment. Briefly, ion transfer tube temp = 300 °C, Easy-IC internal mass calibration, default charge state = 2 and cycle time = 3 s. Detector type set to Orbitrap, with 60K resolution, with wide quad isolation, mass range = normal, scan range = 300-1500 m/z, max injection time = 50 ms, AGC target = 200,000, microscans = 1, S-lens RF level = 60, without source fragmentation, and datatype = positive and centroid. MIPS was set as on, included charge states = 2-6 (reject unassigned). Dynamic exclusion enabled with n = 1 for 30 s and 45 s exclusion duration at 10 ppm for high and low. Precursor selection decision = most intense, top 20, isolation window = 1.6, scan range = auto normal, first mass = 110, collision energy 30%, CID, Detector type = ion trap, OT resolution = 30K, IT scan rate = rapid, max injection time = 75 ms, AGC target = 10,000, Q = 0.25, inject ions for all available parallelizable time.

### MS data analysis and quantification

Protein identification/quantification and analysis were performed with Integrated Proteomics Pipeline - IP2 (Bruker, Madison, WI. http://www.integratedproteomics.com/) using ProLuCID^64^, DTASelect2^65^, Census and Quantitative Analysis. Spectrum raw files were extracted into MS1, MS2 files using RawConverter (http://fields.scripps.edu/downloads.php). The tandem mass spectra (raw files from the same sample were searched together) were searched against UniProt mouse (downloaded on 07-29-2023) protein databases ^66^ and matched to sequences using the ProLuCID/SEQUEST algorithm (ProLuCID version 3.1) with 50 ppm peptide mass tolerance for precursor ions and 600 ppm for fragment ions. The search space included all fully and half-tryptic peptide candidates within the mass tolerance window with no-miscleavage constraint, assembled, and filtered with DTASelect2 through IP2. To estimate protein probabilities and false-discovery rates (FDR) accurately, we used a target/decoy database containing the reversed sequences of all the proteins appended to the target database^66^. Each protein identified was required to have a minimum of one peptide of minimal length of six amino acid residues. After the peptide/spectrum matches were filtered, we estimated that the peptide FDRs were ≤ 1% for each sample analysis. Resulting protein lists include subset proteins to allow for consideration of all possible protein isoforms implicated by at least three given peptides identified from the complex protein mixtures. Static modification: 57.02146 C for carbamidomethylation. Differential modification: 42.0106 for acetylation at N-terminals. Quantification was performed by the built-in module in IP2. The raw mass spectrometry data presented in this study was deposited in Mass Spectrometry Interactive Virtual Enviroment (MassIVE) under the identifier MSV000095320 and ProteomeXchange under the identifier PXD053922.

### Software

Spyder (MIT, Python 3.7) was used for MS data analyses. RStudio (version 1.2.1335) was used for MS data visualization. The Database for Annotation, Visualization and Integrated Discovery (DAVID) (https://david.ncifcrf.gov/) was utilized for protein functional annotation analysis. GeneMania (https://genemania.org/) was employed for generating protein-protein interaction networks. Cytoscape (version 3.9.1) was used for constructing and visualizing the networks.

### Statistical Analysis

We used Microsoft Office Excel and GraphPad Prism version 9.0 or 9.3.1 to analyze data and plot graphs. Data were expressed as mean ± s.e.m. For comparisons among 3 groups, one-way ANOVA with Tukey’s multiple comparison test was used; and for comparisons between 2 groups, two-tailed, unpaired Student *t*-tests were used. P ≤ 0.05 was considered significant.

## Acknowledgements

J.S. is supported by the Wellcome Trust, United2EndMND, ARUK/Dementia Consortium, and the Alan Davidson Foundation; J.C., I.B. and M.J.L. are supported by the National Research Foundation of Korea (RS-2023-00261784); J.N.S., Y-Z.W. and K.K.G are supported by the US Public Health Service (AG078796 and S10OD032464), R.G.K., C.D., A.H. and X.G. are supported by the US Public Health Service (NS122908, NS124802), the US Department of Defense (W81XWH-21-1-0236), the Les Turner ALS Foundation and the Heather Koster Family Charitable Fund. We thank Erin Baker and Han-Xiong Deng for technical assistance, critical discussions and project guidance during the undertaking of this work.

