## Supplementary figures and images for "Reduction of RAD23A extends lifespan and mitigates pathology in TDP-43 mice"

### S1

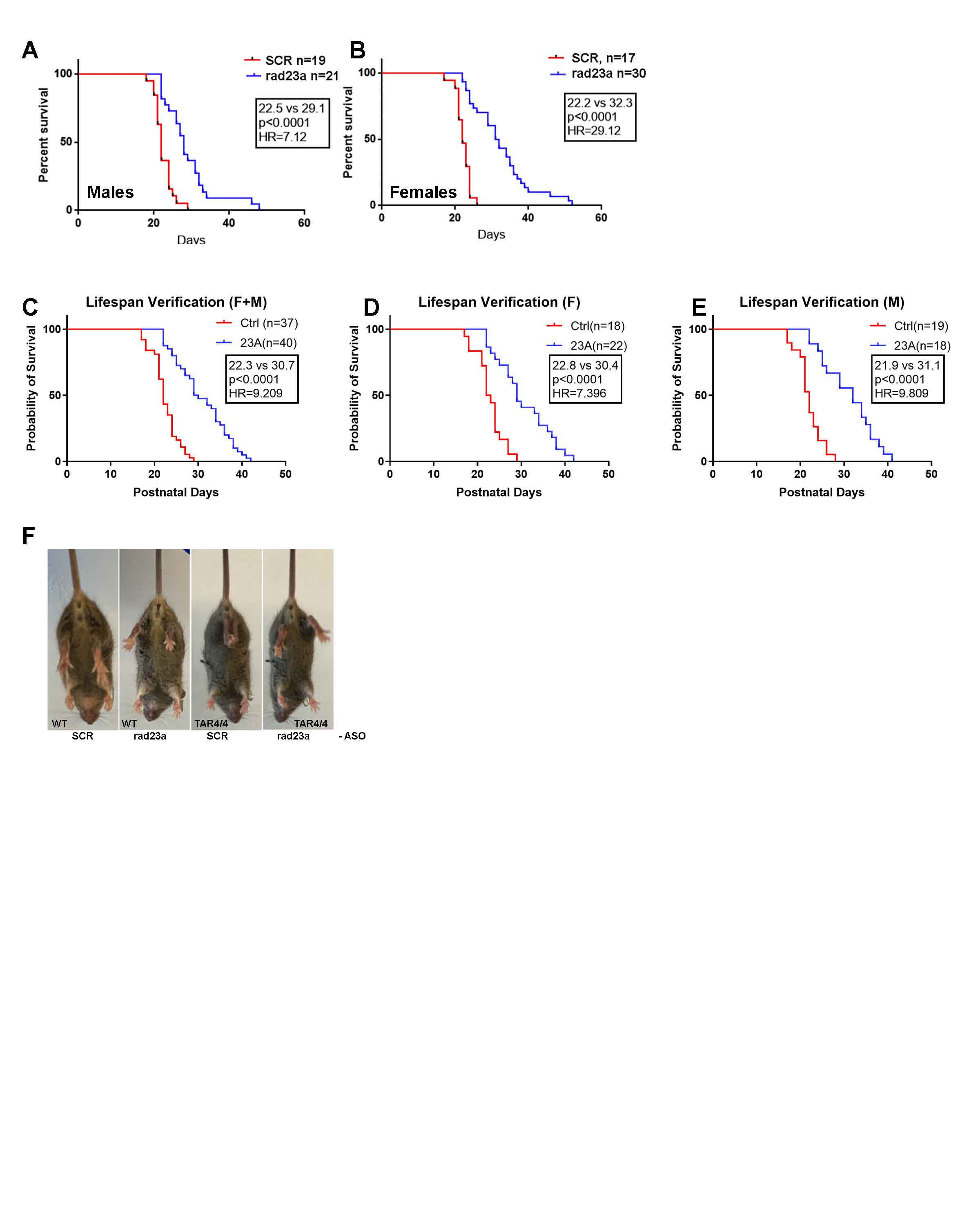

### S6

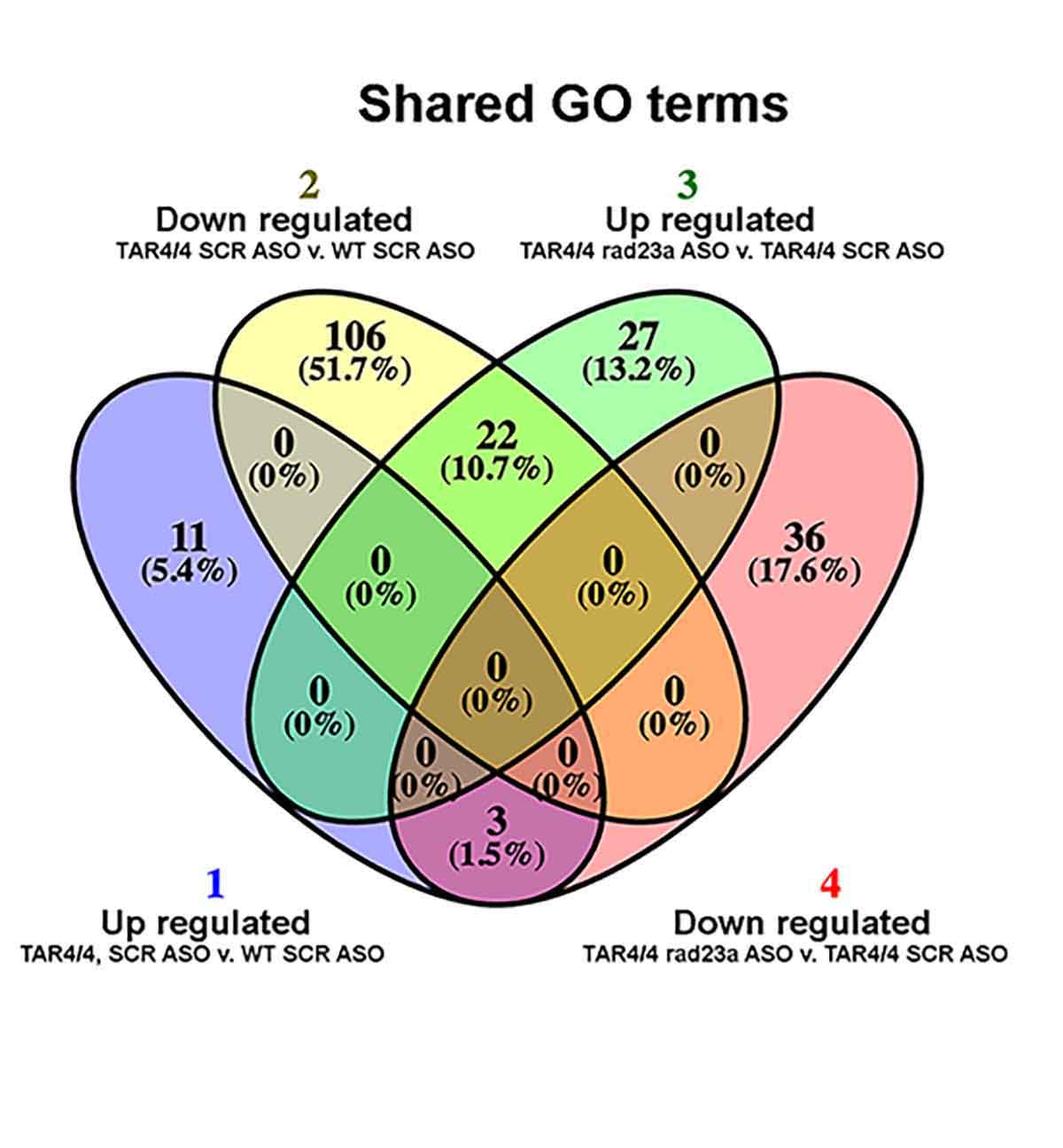

### S7

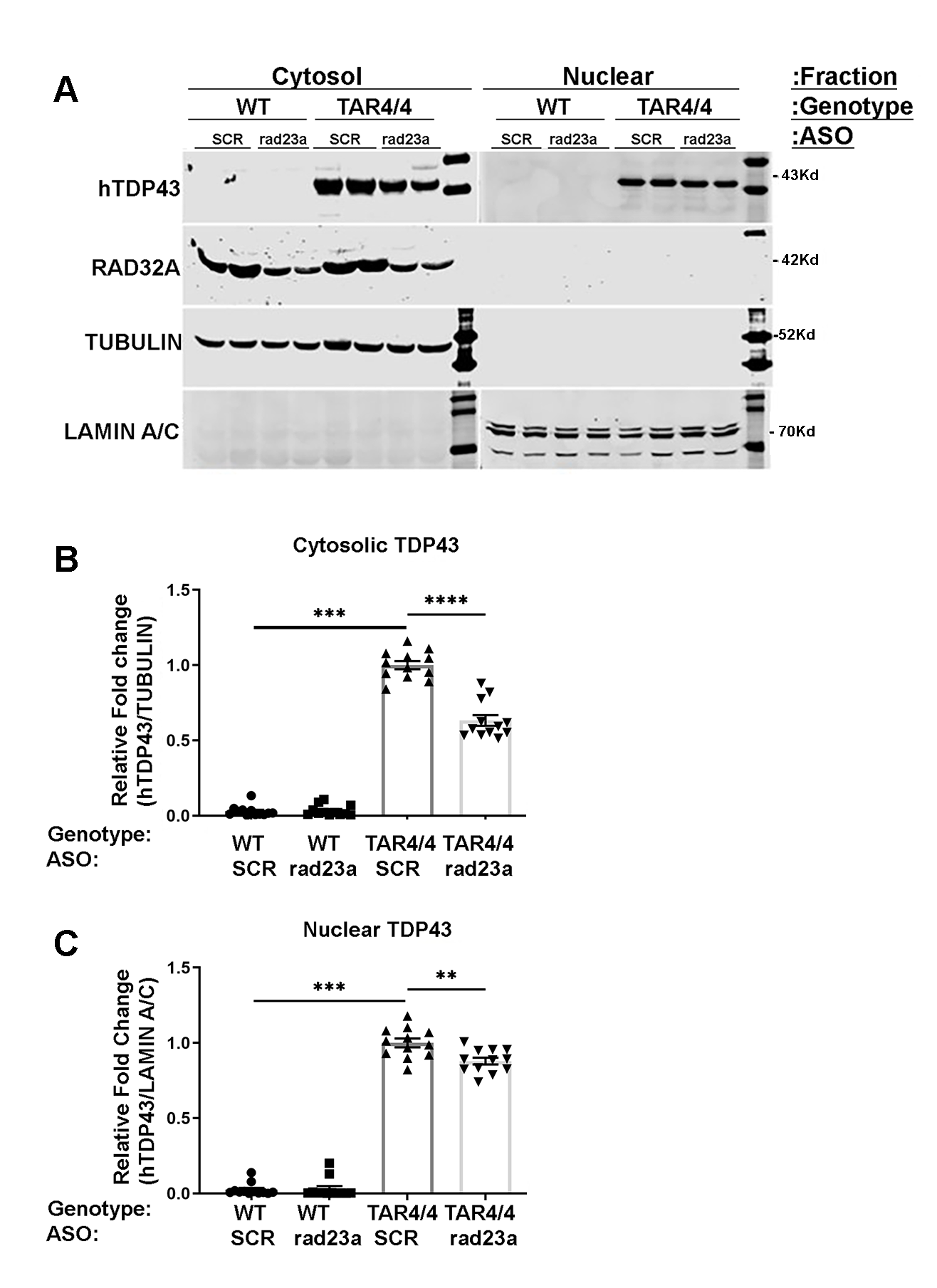

### S8

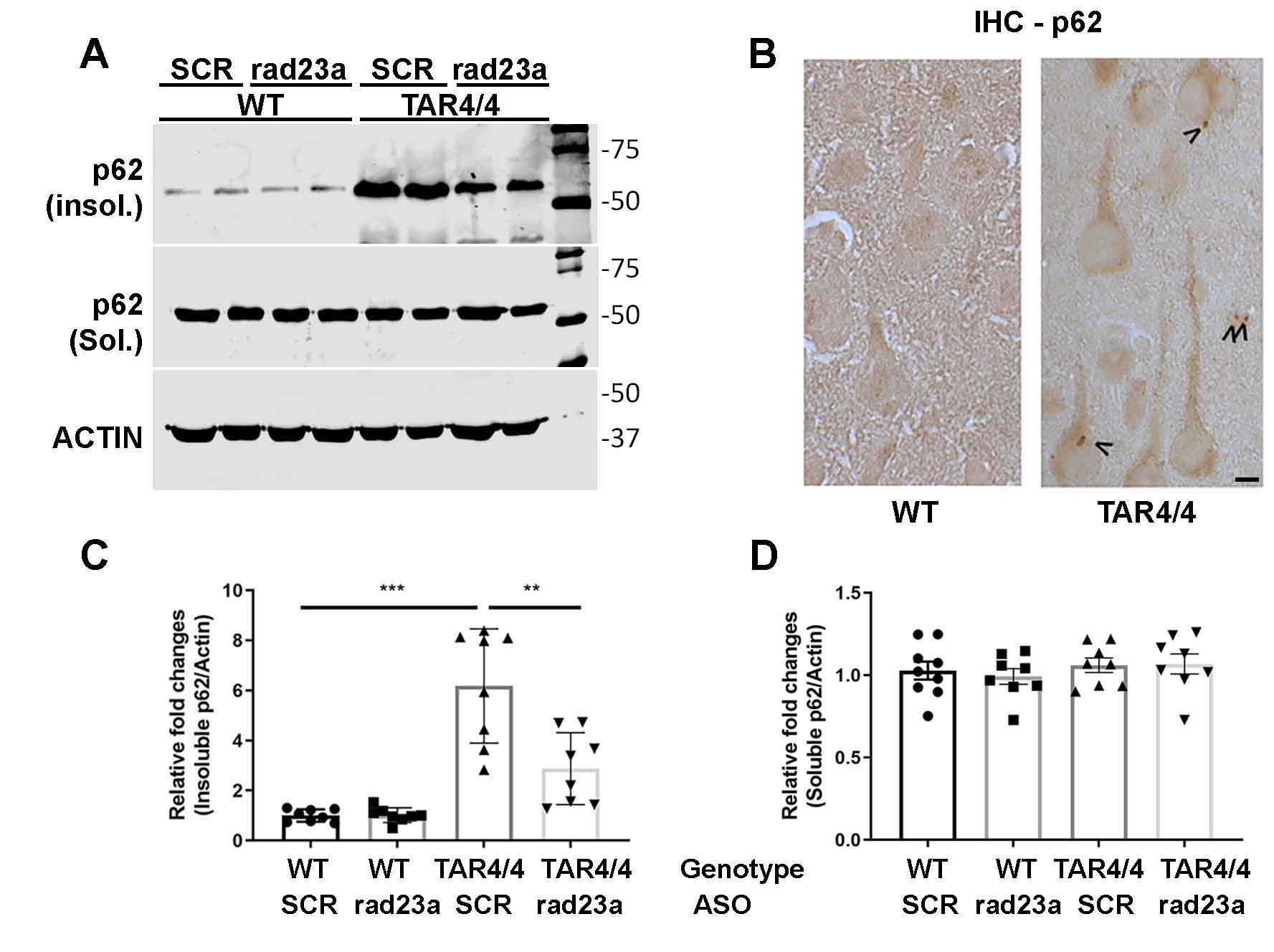

### S9

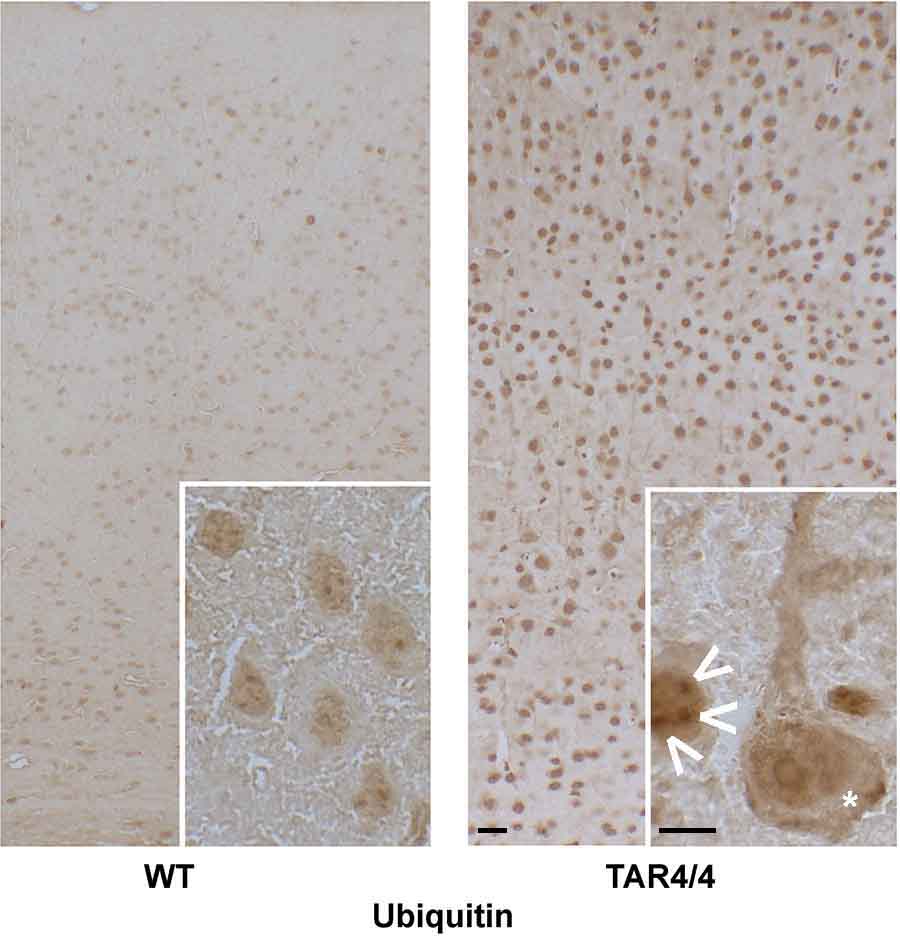

### S11

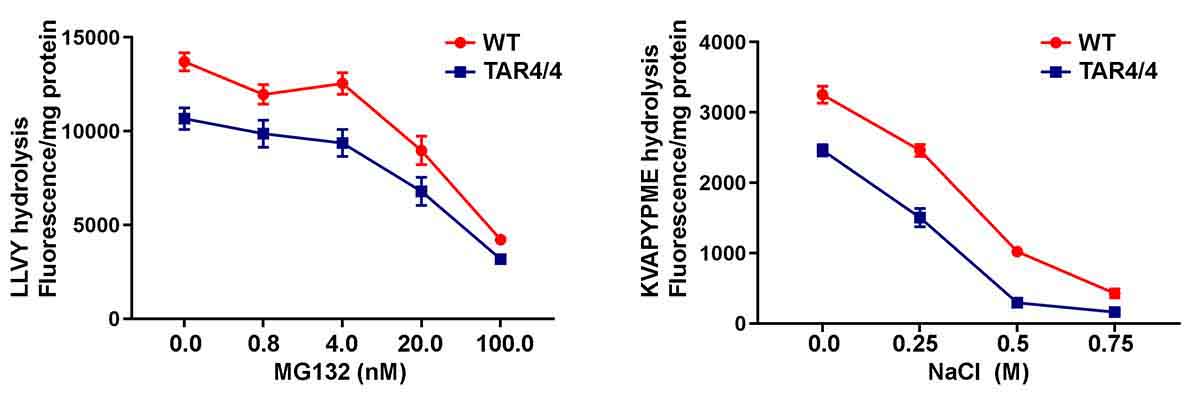

### S12

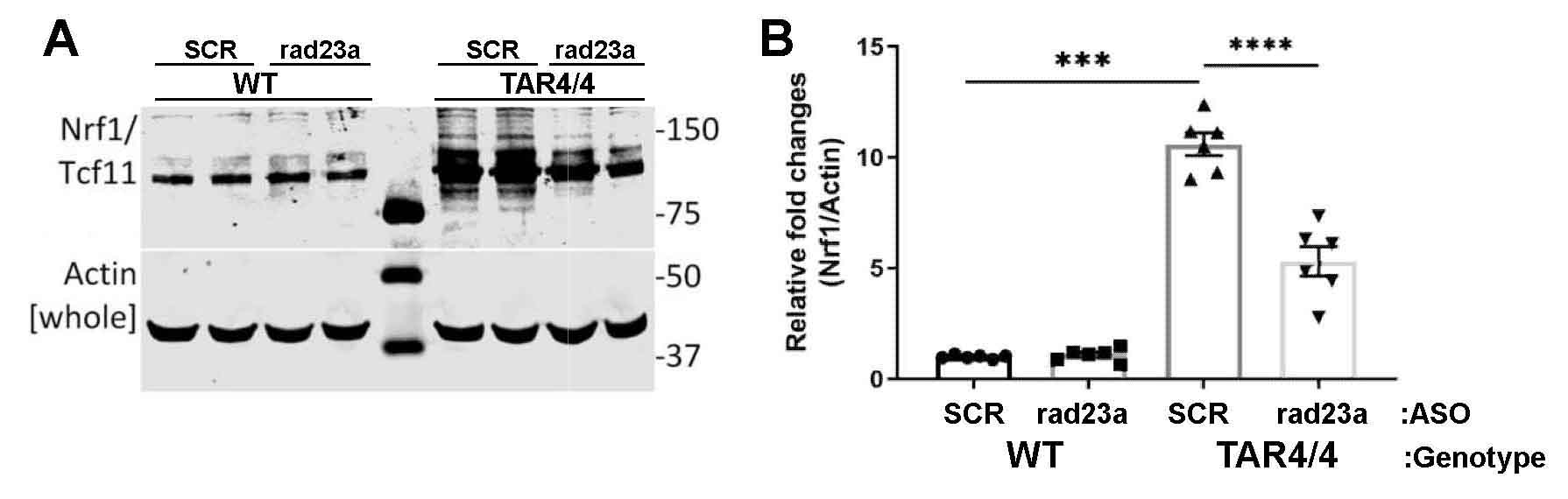
