## Supplementary material for "Reduction of RAD23A extends lifespan and mitigates pathology in TDP-43 mice": S4

**Supplemental Figure 4** Comparison of selected gene expression as determined by RNA-Seq and RT-qPCR. Each group has 4 motor cortex samples for verification with qPCR.

| **Ensembl ID** | **Gene Symbol** | **Fold changes** | | | | |  |  |  |
| --- | --- | --- | --- | --- | --- | --- | --- | --- | --- |
|  |  | **RNA-Seq** | |  | **RT-qPCR** | |  | **T-test** | |
|  |  | Tar4/4_srm vs WT_Srm | Tar4/4_23a vs Tar4/4_Srm |  | Tar4/4_srm vs WT_Srm | Tar4/4_23a vs Tar4/4_Srm |  | Tar4/4_srm vs WT_Srm | Tar4/4_23a vs Tar4/4_Srm |
| ENSMUSG00000004328 | Hif3a | 4.03 | 0.47 |  | 5.13 | 0.64 |  | 0.0121 | 0.1707 |
| ENSMUSG00000021091 | Serpina3n | 5.44 | 0.45 |  | 8.10 | 0.49 |  | 0.0033 | 0.0244 |
| ENSMUSG00000022996 | Wnt10b | 9.69 | 0.27 |  | 8.91 | 0.22 |  | 0.0085 | 0.0215 |
| ENSMUSG00000028364 | Tnc | 0.38 | 3.17 |  | 0.35 | 1.86 |  | 0.0006 | 0.0036 |
| ENSMUSG00000024793 | Tnfrsf25 | 0.35 | 2.99 |  | 0.45 | 3.58 |  | 0.0203 | 0.0098 |
| ENSMUSG00000004891 | Nes | 0.49 | 2.21 |  | 0.58 | 1.57 |  | 0.0522 | 0.0081 |
| ENSMUSG00000074575 | Kcng1 | 0.45 | 2.09 |  | 0.44 | 2.25 |  | 0.0038 | 0.1104 |
| ENSMUSG00000046470 | Sox18 | 0.55 | 2.11 |  | 0.58 | 1.90 |  | 0.0502 | 0.0141 |
| ENSMUSG00000011256 | Adam19 | 0.61 | 2.01 |  | 0.46 | 1.75 |  | 0.0282 | 0.0779 |
| ENSMUSG00000002831 | Plin4 | 24.07 | 0.16 |  | 30.93 | 0.14 |  | 0.0314 | 0.0411 |
| ENSMUSG00000019232 | Etnppl | 7.76 | 0.26 |  | 25.71 | 0.17 |  | 0.0090 | 0.0145 |
| ENSMUSG00000002289 | Angptl4 | 4.12 | 0.33 |  | 6.65 | 0.43 |  | 0.0414 | 0.1343 |
| ENSMUSG00000038393 | Txnip | 2.76 | 0.35 |  | 4.45 | 0.48 |  | 0.0188 | 0.1185 |
| ENSMUSG00000015090 | Ptgds | 1.29 | 0.35 |  | 1.86 | 0.36 |  | 0.0270 | 0.0467 |
| ENSMUSG00000022055 | Nefl | 0.68 | 1.39 |  | 0.49 | 1.84 |  | 0.0794 | 0.0343 |

D

**Table Sn’** Primers used in the RT-qPCR.

| **Primer Name** | **Primer Sequences** | **Primer size (bp)** | **Amplicon size (bp)** |
| --- | --- | --- | --- |
| Hif3a-Fwd | CCCAGTCGGAAAGTATCATCTG | 22 | 94 |
| Hif3a-Rev | GAGTATGTTGCTCCGTTTGTTC | 22 |  |
| Serpina3n-Fwd | TCTCCATCTCCACCGACTAC | 20 | 110 |
| Serpina3n-Rev | AGATCCTTGGTTCCTGTGATTG | 22 |  |
| Wnt10b-Fwd | TCTCTCGGGATTTCTTGGATTC | 22 | 121 |
| Wnt10b-Rev | CATTTGCACTTCCGCTTCAG | 20 |  |
| Tnc-Fwd | TGATACTACCTCTGGCCTCTAC | 21 | 104 |
| Tnc-Rev | AACAATCCATCCACCTCCATC | 22 |  |
| Tnfrsf25-Fwd | CAGCTCTACGATGTGATGGATG | 22 | 99 |
| Tnfrsf25-Rev | ACCTCCACGGCTTCAATTTC | 20 |  |
| Nes-Fwd | GAAGCACTGGGAAGAGTAGAAG | 22 | 110 |
| Nes-Rev | CTAACTCATCTGCCTCACTGTC | 22 |  |
| Robo3-Fwd | GCAGTCCACAGCTACTCTTAC | 21 | 124 |
| Robo3-Rev | GATACGCTGAGAGGAGAACTTG | 22 |  |
| Kcng1-Fwd | CCATACTTTCTCCCGCTCTTAC | 22 | 140 |
| Kcng1-Rev | GGCACTTCCAAACAAGATGTC | 21 |  |
| Sox18-Fwd | CAATCCTCTGTCACCACCAC | 20 | 126 |
| Sox18-Rev | TCCGGCTGCAATTGAGATAC | 20 |  |
| Adam19-Fwd | TACAGACAGAGCCACAAACTG | 21 | 101 |
| Adam19-Rev | CAGTTCCACCACTCTGAGATAC | 22 |  |
| Plin4-Fwd | TGTGTCTGGTATGGCTTCATC | 21 | 121 |
| Plin4-Rev | AAGATATCACCCAGCCCTTTC | 21 |  |
| Etnppl-Fwd | AGCTGCTGAGTGAACAGAAG | 20 | 95 |
| Etnppl-Rev | TCACGGTCCTTCACCAAATC | 20 |  |
| Angptl4-Fwd | GGACCTTAACTGTGCCAAGAG | 21 | 97 |
| Angptl4-Rev | CGTGGGATAGAGTGGAAGTATTG | 23 |  |
| Txnip-Fwd | CTTCGAAGTGATGGATCTAGTGG | 23 | 99 |
| Txnip-Rev | CAGGAATGAACATGCAGGAAAC | 22 |  |
| Ptgds-Fwd | GACGAGTACGCTCTGCTATTC | 21 | 122 |
| Ptgds-Rev | AGGTGGTGAATTTCTCCTTCAG | 22 |  |
| Nefl-Fwd | GCCTTGGACATCGAGATTGCAG | 22 | 141 |
| Nefl-Rev | CAAGCCACTGTAAGCAGAACGG | 22 |  |
